# Color, activity period, and eye structure in four lineages of ants: pale, nocturnal species have evolved larger eyes and larger facets than their dark, diurnal congeners

**DOI:** 10.1101/2021.09.13.460014

**Authors:** Robert A. Johnson, Ronald L. Rutowski

**Affiliations:** School of Life Sciences Arizona State University Tempe, AZ 85287-4501, USA

## Abstract

The eyes of insects display an incredible diversity of adaptations to enhance vision across the gamut of light levels that they experience. One commonly studied contrast is the difference in eye structure between nocturnal and diurnal species, with nocturnal species typically having features that enhance eye sensitivity such as larger eyes, larger eye facets, and larger ocelli. In this study, we compared eye structure between workers of closely related nocturnal and diurnal above ground foraging ant species (family Formicidae) in four genera (Myrmecocystus, Aphaenogaster, Temnothorax, Veromessor). In all four genera, nocturnal species tend to have little cuticular pigment (pale), while diurnal species are fully pigmented (dark), hence we could use cuticle coloration as a surrogate for activity pattern. Across three genera (Myrmecocystus, Aphaenogaster, Temnothorax), pale species, as expected for nocturnally active animals had larger eyes, larger facet diameters, and larger visual spans compared to their dark, more day active congeners. This same pattern occurred for one pale species of Veromessor, but not the other. There were no consistent differences between nocturnal and diurnal species in interommatidial angles and eye parameters both within and among genera. Hence, the evolution of eye features that enhance sensitivity in low light levels do not appear to have consistent correlated effects on features related to visual acuity. A survey across several additional ant genera found numerous other pale species with enlarged eyes, suggesting these traits evolved multiple times within and across genera. We also compared size of the anterior ocellus in workers of pale versus dark species of Myrmecocystus. In species with larger workers, the anterior ocellus was smaller in pale than in dark species, but this difference mostly disappeared for species with smaller workers. Presence of the anterior ocellus also was size-dependent in the two largest pale species.

## Introduction

Ectotherms display extensive variation in color that arises at least in part from variation in the amount of pigment deposited in the cuticle/integument, with melanin being the most common pigment [1, 2]. Diverse selective factors favor the evolution of dark body coloration including biotic factors such as predation and sexual selection [3–6], and abiotic factors such as temperature, ultraviolet–B radiation, and desiccation [7–11]. Despite the various potential benefits of melanin deposition, numerous clades contain species with little or no pigment in their integument. These pale species occur in various taxa including fish, salamanders, insects, shrimp, and spiders [12–15].

One common correlate of pigment level in the integument is light environment. Pale animals with little to no melanin are common in environments with no ambient light such as caves, the deep sea, soil, and parasites inside the body of hosts [14, 16, 17], but are rare in terrestrial environments. Animals that live in dim light conditions, i.e., active nocturnally, sometimes also have little pigmentation and thus are pale compared to their diurnal congeners [e.g., bees, 18, 19]. However, the adaptive advantage of low pigment levels in low-light environments is unclear given that melanin is relatively cheap to produce and its potential advantages are many [20, 21, but see 22].

Species that deposit little pigment in their integument often display a suite of associated and selectively advantageous traits. One common adaptation in these species, especially those in lightless environments, is a severe reduction in or loss of eyes, with this trait being particularly well-studied in fish and other species that have pigmented terrestrial counterparts with fully developed eyes [12, 13, 23, 24]. Alternatively, many organisms that live in dim light environments and have lost some to most of their pigment possess exceptionally large eyes that enhance visual system performance in low light [18, 19].

Herein, we explore the association between body coloration, daily activity patterns, and eye structure in ant species that vary in the extent of their cuticular pigmentation. We designate two categories of coloration: pale (little pigmentation, appearing mostly concolorous whitish-yellow to yellowish to amber), and dark (extensive pigmentation, appearing orange or light to dark brown or black) (see Figs 1-4). Existing knowledge about the relationship between body color, light environment, and eye structure in ants suggests that they display relationships common in other taxa, i.e., (1) compared to close relatives from well-lit environments, species that live in lightless subterranean habitats are paler in color and have eyes that are absent or severely reduced in size [e.g., 25, 26], and (2) pale, nocturnal species that forage above ground have relatively large eyes compared to diurnally foraging species [e.g., 27]. Specifically, this study was motivated by the observation that eyes and facet lenses were larger in pale, nocturnal, above ground foraging species of honey pot ants (*Myrmecocystus* subgenus *Myrmecocytus* – subfamily Formicinae) compared to their dark congeners [see 28].

Broadly, we were interested in the evolution and consequences of these associations, and we used a comparative approach to examine these relationships in four ant genera in two subfamilies (*Myrmecocystus* - subfamily Formicinae and *Aphaenogaster*, *Temnothorax*, *Veromessor* - subfamily Myrmicinae) that contain pale and dark species. This multitaxa approach strengthened our ability to make evolutionary inferences.

We first quantified for each genus the association between cuticular pigmentation and daily activity patterns, i.e., whether pale species are more nocturnal than their dark congeners. We then examined how eye size varies with body color and activity time. Specifically, we determined if within each genus pale species have larger eyes than their dark congeners, i.e., eyes that would enhance vision in dim conditions. Several studies have compared the compound eyes of nocturnal and diurnal ants [27]. However, most of these comparative studies lacked adequate controls for phylogeny in that they compared relatively small numbers of species from different lineages or with unknown phylogenetic relationships [27, 29, 30].

For a subset of these species, we examined eye structure in more detail to explore the effects of activity-pattern-related variation in eye size on visual sensitivity, acuity, and field dimensions. High visual acuity is generally expected for members of the Hymenoptera (i.e., ants, wasps, bees) whose apposition eyes [31] are typically structured to maximize image resolution rather than light capture. Visual resolution of apposition eyes can be assessed by measuring facet diameter (*D*) and interommatidial angle (Δ*ϕ*, the angle between the optical axes of adjacent ommatidia). Larger facets capture more light and *D* is positively correlated with sensitivity, while resolving power is negatively correlated with Δ*ϕ*. Consequently, there may be a potential tradeoff between resolution and sensitivity given that *D,* Δ*ϕ*, and eye size interact in complex and sometimes opposing ways [32]. Nocturnal animals usually resolve this tradeoff in favor of sensitivity, and thus have lower acuity compared to their diurnal counterparts [33]. Hence, we expected *D* to be greater and Δ*ϕ* to be larger in pale species of ants.

We also used the eye parameter (*ρ*, which is the product of *D* in um and Δ*ϕ* in radians) [33–35], to characterize the compromise between sensitivity and acuity for each species. The calculated *ρ* indicates how closely the eye is constructed to the limits imposed by diffraction, i.e., whether the eye is structured to enhance resolution over sensitivity. These values are low for diurnally active species, and they increase at lower light intensities with nocturnal species often having *ρ* values ≥ 2.

For two species within each genus (one pale, one dark), we also measured Δ*ϕ* in the center of the eye, visual field span, and regional variation in *D*. Collectively, these measures permit inferences about how visual field structure varies between nocturnal and diurnal species both within and across genera.

We also examined variation in size of the ocelli in *Myrmecocystus*, which was the only examined genus in which workers possess these structures. Ocelli are a second visual system present in most flying insects that detect polarized light and assist in head stabilization and horizon detection [36–38], but also reflect the natural history and environment of the species [39]. Ocelli are present in alate queens and males of nearly all ant species, but they typically are lacking in the pedestrian workers with *Myrmecocystus* being a notable exception [40, 41]. Snelling [28] noted that the ocelli were smaller in pale than in dark species of *Myrmecocystus*.

All species examined herein were geographically restricted to the southwestern United States and northwestern Mexico. The relatively large number of pale species found in this region suggests that pale ants may occur in other regions of the world and in other genera and/or subfamilies. Consequently, we assessed variation in ant color and its correlation with eye size on a larger geographical and taxonomic scale by surveying photographs across species in several ant genera. Here again, we expected pale body color to be correlated with nocturnal activity as well as eye morphology that enhances visual sensitivity.

## Methods

### Study species

We examined the relationship between body color, activity pattern, and eye structure in 23 ant species spread across four genera in two subfamilies – *Myrmecocystus* (subfamily Formicinae) and *Aphaenogaster*, *Temnothorax*, and *Veromessor* (subfamily Myrmicinae); all four genera contained pale and dark species. Hereafter, names of pale species are in normal font, and names of dark species in **bold** font.

*Myrmecocystus*: We examined 74 workers from six species (Fig 1). This genus is restricted to North America, and it consists of 29 described species [28, 42], plus several undescribed and cryptic species [43]. We compared three size-similar species pairs that differed in pigmentation: small (*M. christineae* and ***M. yuma***), medium (*M. navajo* and ***M. kennedyi***), and large (*M. mexicanus*-02 and ***M. mendax*-03**). All pale species of *Myrmecocystus* occur in two clades, while dark species comprise the rest of the clades in the genus [43].

**Fig. 1.**
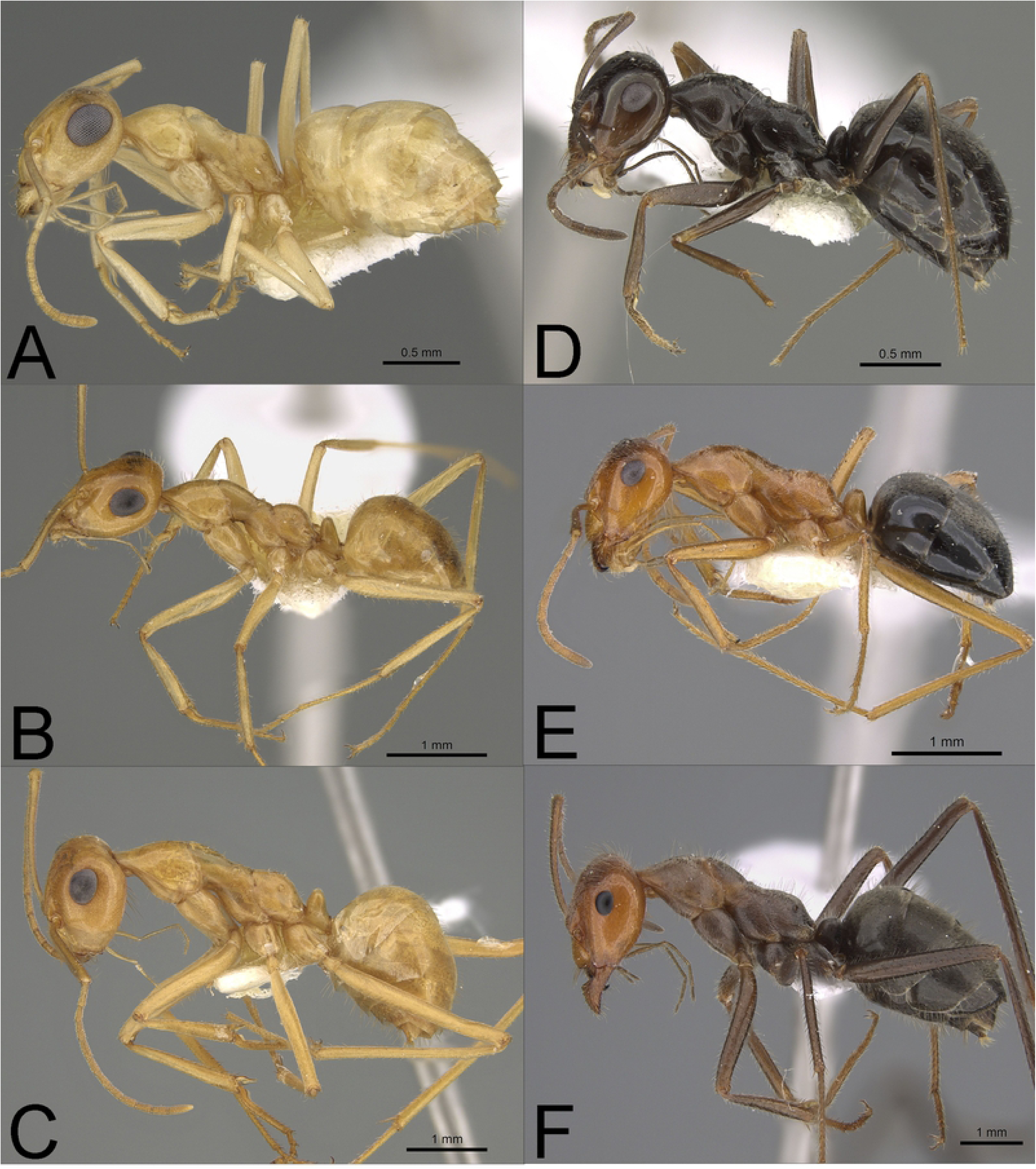
Profile pho togr aphs of pale (A-C) and dark species (D-F) (see text) of *Myrmecocystus* (subfamily Formicinae) examined in this study: (A) *M. christineae* (CAS ENT09233 58), (B) *M. navajo* (CASENT0923356), (C) *M. mexicanus-*02 (CASE NT0923355), (D) *M yuma* (CAS ENT0923359), (E) *M kennedyi* (CASE NT0923362), and (F) *M. mendax-03* (ASUSIBROOOOI 132). Note the relatively larger eyes of pale compared to dark species. Species arearranged by size pairs - A&D, B&E, and C&F (see text). Photographs by Michele Esposito from www.antweb.org.

In his revision of *Myrmecocystus*, Snelling [28] also indicated that the ocelli were reduced in size or absent in pale species compared to their dark congeners. We tested this observation by measuring diameter of the anterior ocellus for workers of the above six species, plus workers of three additional pale species (*M. ewarti*, *M. testaceus*, *M. mexicanus*-01). As such, our analysis included all known pale species except *M. melanoticus* and *M. pyramicus* [28, 43], for which specimens were not available.

*Aphaenogaster*: We examined 38 workers from four species (Fig 2). This genus includes 30 species in North America. We compared *A. megommata*, the only pale species, with three closely related dark congeners ***A. boulderensis***, ***A. occidentalis***, and ***A. patruelis*** [44, 45].

**Fig. 2.**
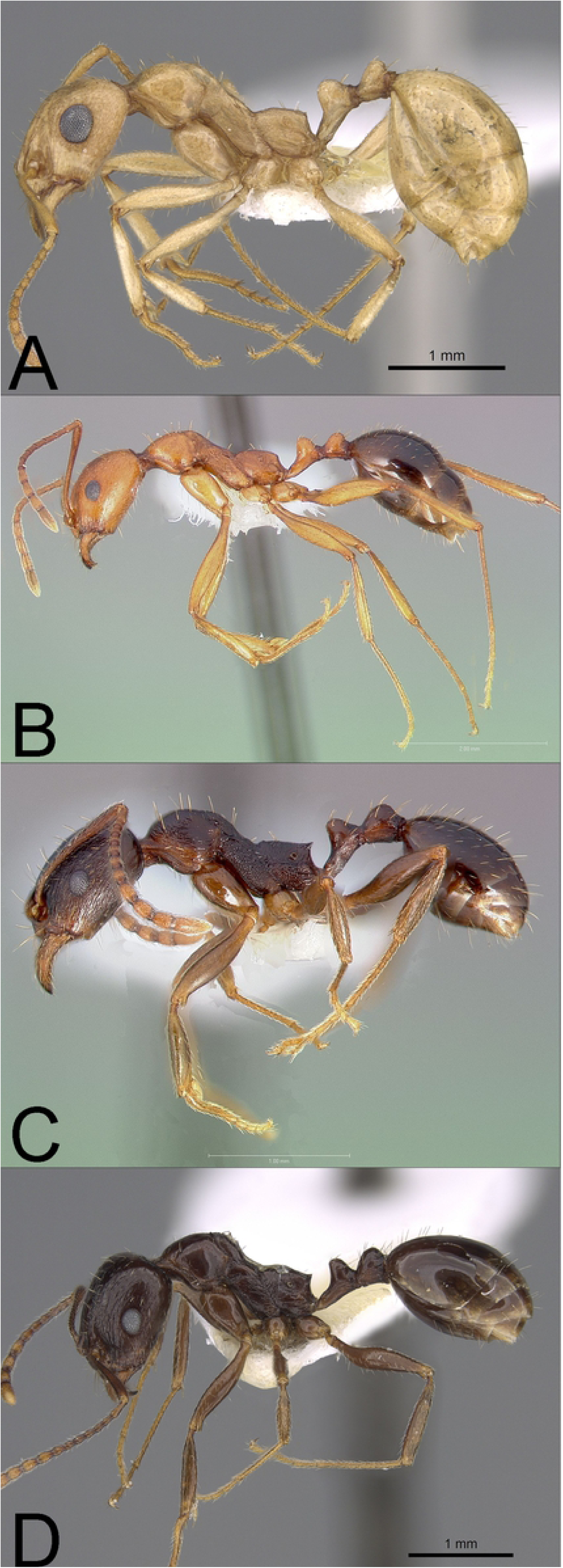
Profile photo graph s of pale (A) and dark species (B-D) (see text) of *Aphaenogaster* (subfamily Myrmicinae) examined in this study : **(A)** *A. megommata* (CASE NT0923367), **(B)** *A. boulderensis* (CASENT0005722), **(C)** *A. occidentalis* (CASE NT0005725), and **(D)** *A. patruelis* (CASE NT0923366). Note the relatively larger eyesof pale compared to dark species. Photographs by Michele Esposito from www.antweb.org.

*Temnothorax*: We examined 29 workers from three species (Fig 3). In North America, this genus consists of more than 80 described species plus numerous undescribed species, with the *T. silvestrii* clade consisting of several poorly known, poorly collected pale species (M. Prebus, pers. comm.). Our analysis included the undescribed pale species *T.* sp. BCA-5 [in 46, as *Leptothorax* sp. BCA-5] from the *T. silvestrii* group, and ***T. neomexicanus*** and ***T. tricarinatus***, which are two dark species from the *T. tricarinatus* group, which is sister to the *T. silvestrii* group [47, M. Prebus, pers. comm.].

**Fig. 3.**
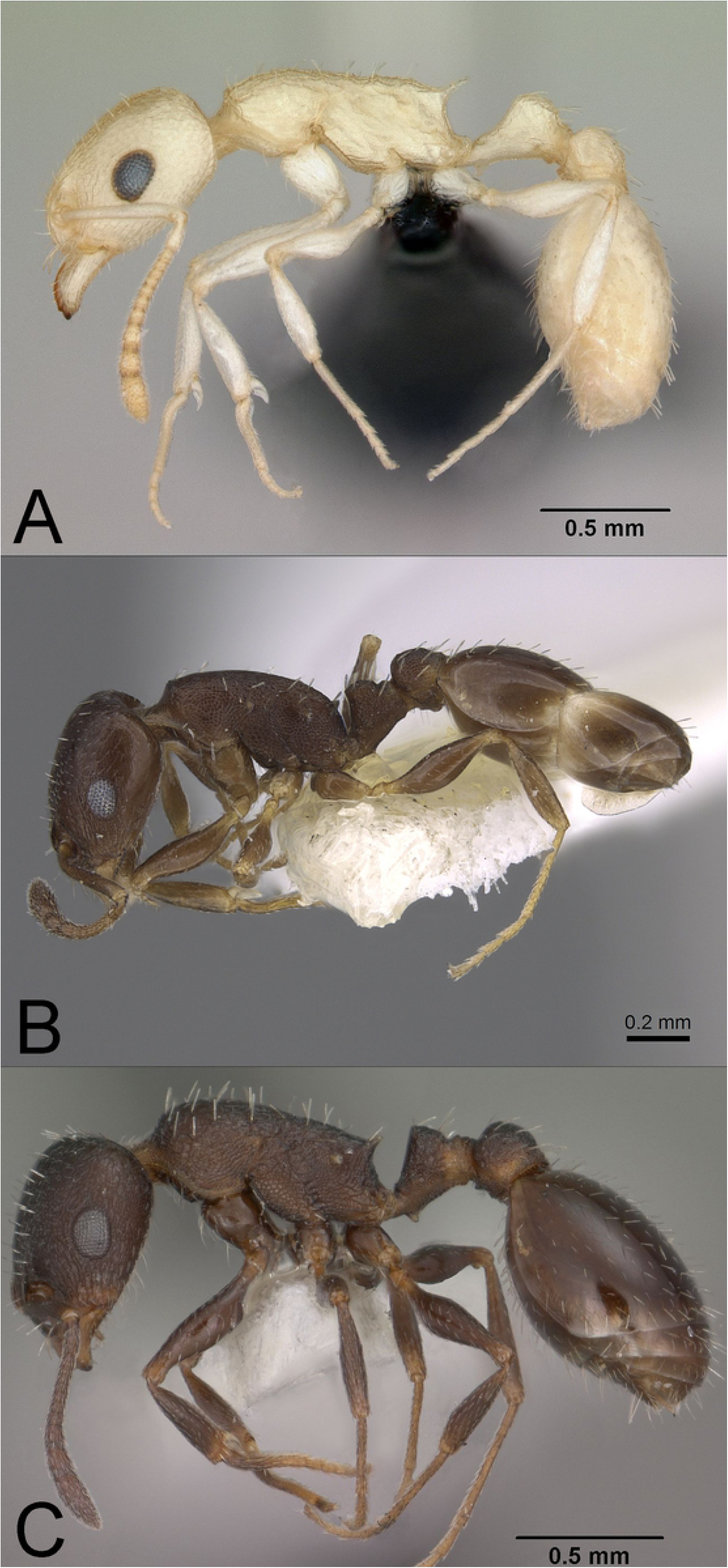
Profile photographs of pale (A) and dark species (B--C) (see text) of *Temnothorax* (subfamilyMyrmicinae) examined in this study: **(A)** *T.* sp. BCA-5 (CASENTOJ 18165), **(B)** *T neomexicanus* (CASENT0923368), and **(C)** *T tricarinatus* (CASENTO I 0 2845) . Note the relatively larger eyes of pale compared to dark species. Photographs by Michele Esposito from www.antweb.org.

*Veromessor*: We examined 133 workers from the 10 species that occur in the genus (Fig 4). This genus only occurs in North America [48, 49]; two species are pale, *V. lariversi* and *V.* RAJ*-pseu*, while the eight other species are dark.

**Fig. 4.**
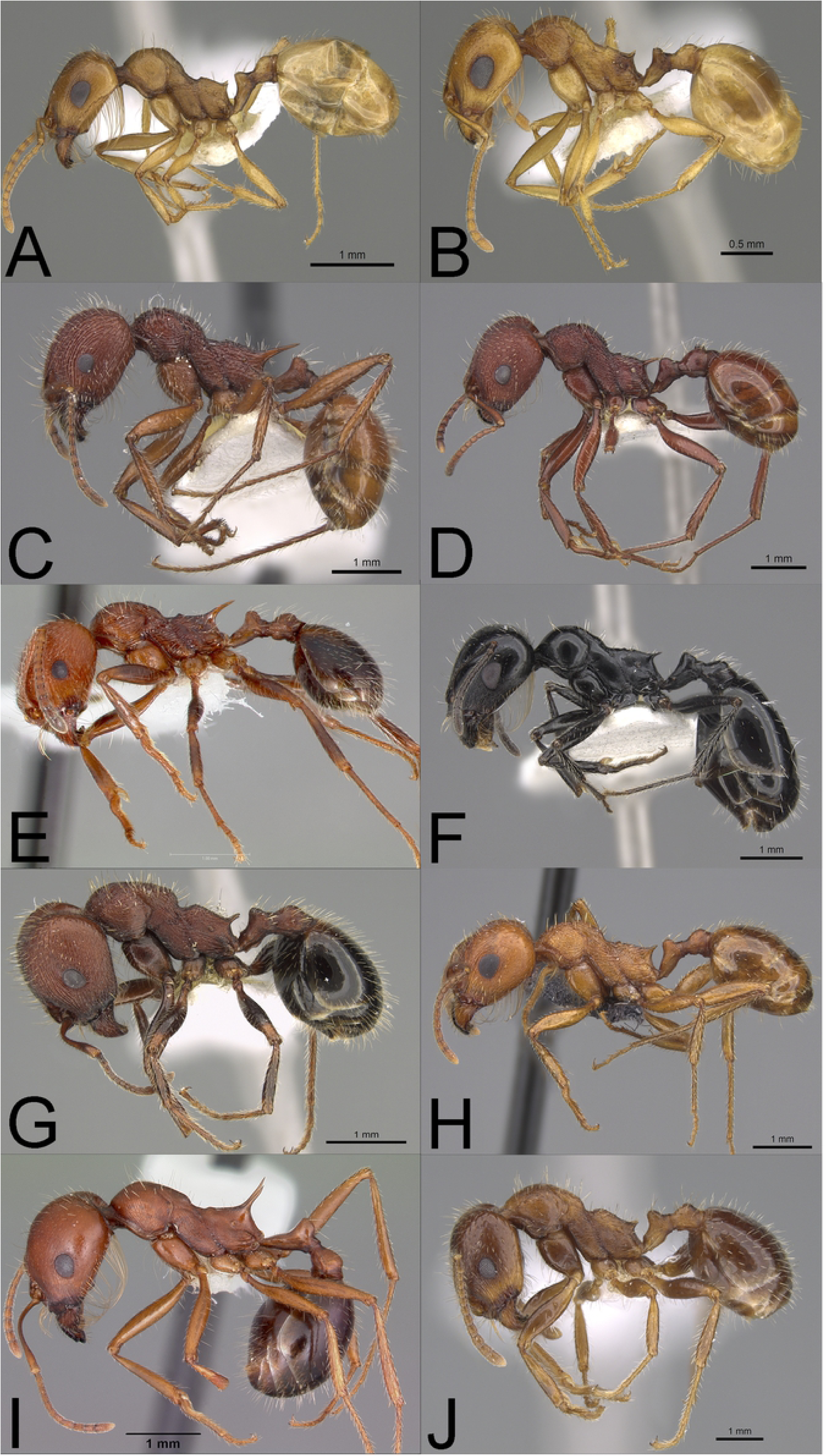
Profile photographsof pale **(A-B)** and dark species **(C-J)** (see text) of *Veromessor* (subfamily Myrmicinae) examined in this study: **(A)** *V. lariversi* (CASENT0923345), **(B)** *V. RAJ-pseu* ( CASENT0923346), **(C)** *V. andrei* (CASENT0923 l37), **(D)** *V. lobognathus* (CASENT0923126), **(E)** *V. chamberlini* (CASENT0005730), **(F)** *V. pergandei* (CASENT0923124), **(G)** *V. chicoensis* (CASENT0923347), **(H)** *V. smithi* (MCZ-ENT00671466), (I) *V.julianus* (CASENTOI 04946), and **(J)** *V. stoddardi* (CASENT0922825). Note the relatively larger eyes of pale compared to dark species. Photographs by Wade Lee, April Nobile, and Michele Esposito from www.antweb.org.

All specimens are deposited in the collections of Robert. A. Johnson, Tempe, AZ (RAJC), Matthew M. Prebus, Tempe, AZ (MMPC), and the Social Insect Biodiversity Repository (SIBR), Arizona State University, Tempe, AZ.

### Measure of body coloration

All brightness, body size, and eye measurements were made from photographs of workers as described below. We quantified worker color using the brightness value (v or B, in HSV format), which is similar to the lightness value in HSL that has been used to characterize body color in other studies of ants [8, 10]. Brightness (B) was measured using the color window in Adobe Photoshop from photographs downloaded from Antweb (www.antweb.org). Obviously discolored specimens were excluded, i.e., those in which the color differed substantially from intraspecific specimens recently collected by RAJ. Using the photograph of the worker body in profile, we measured B on the head (immediately posterior to the eye), mesosoma (center of mesopleura), and gaster (anteroposterior portion of first gastral tergum), then averaged these values for each worker, then averaged that value across all workers for each species. We compared mean B values for pale versus dark species using a t-test.

### Activity patterns

The relationship between color and activity pattern was evaluated by gleaning above-ground foraging times from literature, personal observations, and personal communications. Foraging times were classified as one or more of the following: diurnal, nocturnal, matinal, crepuscular, and variable. The category “variable” included species in which foraging time varies seasonally with temperature – diurnal when days are cool, crepuscular-matinal as temperatures increase, and nocturnal when nights are warm. Exclusively day-active or night-active species were classified as diurnal and nocturnal, respectively. Species that forage during both day and night (e.g., variable) and those that have matinal and crepuscular foraging were classified as variable. We tested the association between color (pale and dark) and activity time (diurnal, nocturnal, variable) using a Fisher’s exact test [50].

### Body size and eye measurements

We measured body size and eye characteristics for workers from all 23 species listed above. Body size was measured as mesosoma length, which is a standard measure for body size in ants. Mesosoma length was measured as the diagonal length of the mesosoma in profile from the point at which the pronotum meets the cervical shield to the posterior base of the metapleural lobe. Mesosoma length was measured from photographs taken using a Spot Insight QE camera attached to a Leica MZ 12_5_ microscope. Images were then displayed on a video monitor, and mesosoma length was measured using ImageJ (available at http://rsb.info.nih.gov/nih-image/). Measurements were calibrated using photographs of an ocular micrometer scaled in 0.01 mm increments.

Eye measurements were made from high-resolution photographs of the left eye taken in profile focused on the center of the eye at an angle that allowed viewing all facets. Photographs were taken using a Leica M205C microscope at 100× that was linked to the stacking software program Helicon Focus (www.heliconsoft.com/heliconsoft-products/helicon-focus). This software combines photographs taken in different focus planes into one photograph where the entire eye surface is in focus. Facet lenses were counted, and eye area and facet diameter (*D*) were measured using Digimizer (https://www.digimizer.com/). The area tool was used to calculate area. This tool also calculated the centroid of the eye, and *D* was the average of four adjoining facets at the centroid. We also measured the diameter of the anterior ocellus for species of *Myrmecocystus*. All photographs contained a 0.15 mm scale bar for calibrating measurements.

### Detailed eye measurements

Detailed measures of eyes and visual field were taken from a subset of the 23 species. We conducted four eye measurements on two species from each genus, one pale, the other dark that were closely matched in mesosoma length.

#### Interommatidial angle (**Δ*ϕ***), eye parameter (***ρ***), and visual field

Measurements of Δ*ϕ* allowed us to examine how spatial resolution varied with activity period, eye size, and body size. We measured Δ*ϕ* in the lateral region of the eye for five workers from one pale and one dark species in each of the four genera (*M. navajo*, ***M. kennedyi*** in *Myrmecocystus*; *A. megommata*, ***A. patruelis*** in *Aphaenogaster*; *T*. BCA-5, ***T. neomexicanus*** in *Temnothorax*; *V. lariversi*, ***V. chicoensis*** in *Veromessor*) using the radius of curvature method outlined in Bergman and Rutowski [51] with minor modifications. For each specimen, we photographed the left eye in side view from a position on a line perpendicular to the anterior-posterior axis of the eye. This created a photograph showing the edge of the eye surface corresponding at its apex in side view with individual facets visible at the eye edge. Each image was copied into Geogebra (©International Geogebra Institute, 2013; www.geogebra.org) to measure the angle subtended by the eye surface spanned by two facets in the apical region. Briefly, a point was identified between two facets at the edge of the eye at its apex. We then drew a line to a point on the eye surface two facet rows away in the anterior direction. This was taken as the tangent to the eye at that point and the perpendicular bisector of these lines between these points was drawn. This was also done for a line extending from the original point to a point two facet rows in the posterior direction. The Δ*ϕ* was the angle between these two perpendicular bisectors divided by two. We measured Δ*ϕ* three times in the same area for each worker and used the average in our analysis. We calculated *ρ* for each worker as the product of Δ*ϕ* in radians and *D* in μm.

The same images that were used to measure Δ*ϕ* also were used to measure the anterior-posterior visual field span. Lines perpendicular to the tangent of the eye surface were drawn at the anterior and posterior edge of each eye. The angular span of the visual field along the anterior-posterior axis was characterized by the angle between these lines. This measurement was repeated three times for each specimen and the mean value was used in our analyses.

#### Regional variation in *D*

Regional variation in *D* was measured for *Myrmecocystus* and *Veromessor*. The small eyes of *Aphaenogaster* and *Temnothorax* contained too few facets to warrant examination of regional variation. We quantified regional variation in *D* using the photographs taken for eye size measurements (see above) for one pale and one dark species of *Myrmecocystus* (*M. navajo* and ***M. kennedyi***) and *Veromessor* (*V. lariversi* and ***V. chicocensis***). The image for each individual was printed on letter size paper, and *D*’s were measured in five eye regions: anterior, dorsal, lateral, posterior, and ventral. The anterior-posterior axis of the eye was a line from the mandible to the posterior corner of the head, and eye regions were described relative to this line. In each region, three facets in one row were measured to the nearest 0.1 mm with digital calipers, then scaled using the 0.15 mm scale bar present on each photograph. Mean *D* in each region was total length divided by three.

### Data analysis

Eye area, facet number, and mean *D* (dependent variables) were compared across species (independent variable) within each genus using a multivariate analysis-of-covariance (MANCOVA) in the general linear models (GLM) program of SPSS [50]; mesosoma length was the covariate. A least significant difference (LSD) post-hoc test compared the estimated marginal means across species for each variable using mesosoma length as a covariate. Diameter of the anterior ocellus (dependent variable) was compared similarly across species (independent variable) of *Myrmecocystus* using analysis-of-covariance (ANCOVA). *Myrmecocystus navajo* was omitted from this analysis because only one worker had a visible anterior ocellus (see below).

Regional variation in *D* was analyzed using a one-way repeated measures ANOVA followed by a LSD post-hoc test for each species [50]. The dependent variables - Δ*ϕ*, *p*, and visual field span - were compared in separate ANCOVA’s using genus (4 levels) and activity period (diurnal versus nocturnal) as independent variables, with mesosoma length as a covariate [50]. A Tukey’s HSD post-hoc test compared differences across genera and species for each variable. For all tests, data were transformed, as necessary, to meet the assumptions for homogeneity of variance (Box’s M test and Levene’s test) and homogeneity of regression slopes.

### Survey for additional pale ant species

We used Antweb (www.antweb.org) to scan photographs for pale ant species in the genera *Aphaenogaster*, *Crematogaster*, *Messor*, and *Temnothorax* (subfamily Myrmicinae), and *Dorymyrmex* and *Iridomyrmex* (subfamily Dolichoderinae). We scrolled through frontal photographs of the head for all species in each of these genera looking for species that appeared pale and that also appeared to have eyes that were larger than those of nearby dark species (e.g., https://www.antweb.org/images.do?subfamily=myrmicinae&genus=temnothorax&rank=genus&project=allantwebants). We verified our visual assessment of color for these taxa by measuring their brightness value (B) using Adobe Photoshop, as detailed above.

## Results

### Pigmentation and daily activity pattern

We first quantitatively confirmed our visual impressions of variation in body color. As predicted, our brightness values (B) measured from Antweb photographs were consistently and significantly higher for species that we visually classified as pale compared to dark across all four genera (t-test, t=-9.8_24 df_, *P* < 0.0001). Mean B values were 76.3 (*n* = 10) for pale species and 42.1 (*n* = 16) for dark species (Table 1, Fig 5). Values did not overlap for any pale and dark species as all dark species had a mean B below 60, whereas all pale species had a mean B above 65. However, note that the two pale species of *Veromessor* displayed mean B values that ranged from 65–70, which was intermediate to pale and dark species in the other three genera (Fig 5). There also was a significant effect of color (pale, dark) on activity period (diurnal, nocturnal, variable) (*P* < 0.0001, two-sided Fisher’s exact test) with a preponderance of pale species that are nocturnal and dark species that have diurnal/variable activity periods (Table 2). Henceforth, we use pale and dark to refer to species with nocturnal and diurnal/variable activity periods, respectively.

**Fig. 5.**
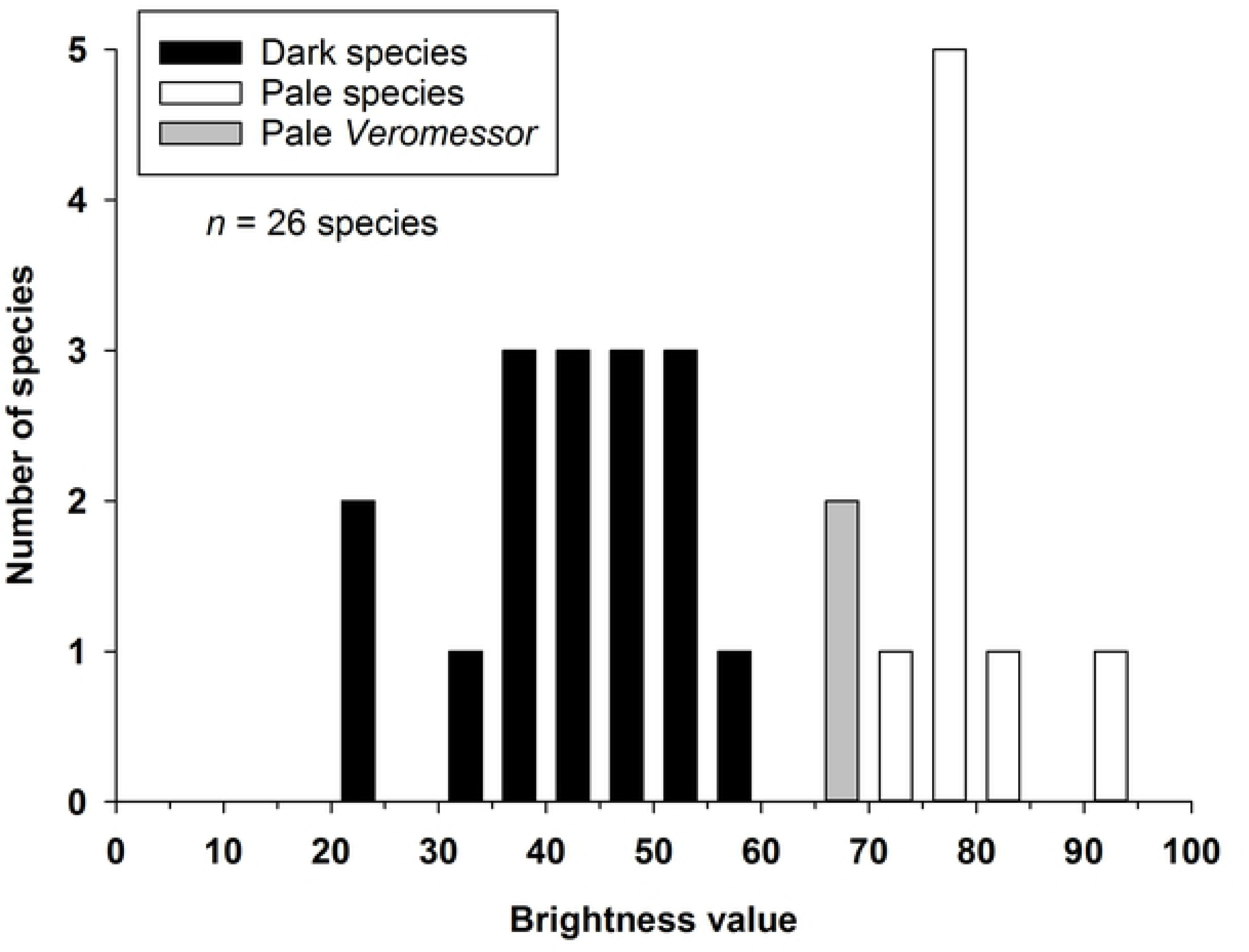
Brightness values for the 26 ant species examined in this study (see text and Table I, Figs 1-4).

**Table 1.** Foraging time for ant species examined in this study. Species are listed alphabetically by subfamily, genus, and species within each genus. Pale species (see text) are in normal font; dark species are in **bold** font. Brightness values (B) are given as mean ± 1 SE (number of workers) (see text). Taxonomy follows Bolton [52]; species followed by a number are undescribed or in the process of revision [see 43].

| Species | Brightness value | Foraging time | References |
| --- | --- | --- | --- |
| **Subfamily Formicidae** |  |  |  |
| ***Myrmecocystus*** |  |  |  |
| *christineae* Snelling | 80.7 ± 2.2 (4) | nocturnal* | [53] |
| *ewarti* Snelling<sup>#</sup> | 74.9 ± 1.7 (3) | nocturnal | [28, 53] |
| ***kennedyi* Snelling** | 51.9 ± 3.7 (3) | diurnal | [28, 53] |
| **sp. mendax-03** | 59.0 ± 3.2 (4) | diurnal | [28] |
| sp. *mexicanus*-01<sup>#</sup> | 77.0 ± 3.2 (4) | nocturnal | [28, 53] |
| sp. *mexicanus*-02 | 76.3 ± 3.7 (3) | nocturnal | [28, 53] |
| *navajo* Wheeler | 76.6 ± 2.4 (6) | nocturnal, crepuscular in cooler months | [28, 53] |
| *testaceus* Emery<sup>#</sup> | 75.2 ± 5.1 (3) | nocturnal | [28, 53] |
| ***yuma* Wheeler** | 35.3 ± 0.8 (3) | matinal-crepuscular | [28] |
| **Subfamily Myrmicinae** |  |  |  |
| ***Aphaenogaster*** |  |  |  |
| *boulderensis* Smith | 46.5 ± 7.2 (2) | crepuscular, nocturnal, matinal | [53] |
| *megommata* Smith | 78.1 ± 4.3 (6) | crepuscular, nocturnal | [53, 54] |
| *occidentalis* (Emery) | 46.1 ± 3.3 (8) | variable<sup>+</sup> | B. DeMarco, pers. comm.; P.S. Ward, pers. comm. |
| *patruelis* Forel | 43.1 ± 5.2 (9) | variable | D. Holway, pers. comm.; P.S. Ward, pers. comm. |
| **Subfamily Myrmicinae** |  |  |  |
| ***Temnothorax*** |  |  |  |
| sp. BCA-5 | 86.8 ± 1.2 (2) | nocturnal | [46, as *Leptothorax* sp. BCA-5]; R.A. Johnson, pers. obs. |
| *neomexicanus* (Wheeler) | 23.3 ± 3.4 (4) | crepuscular | S.P. Cover, pers. comm. |
| *tricarinatus* (Emery) | 32.7 ± 2.5 (3) | crepusclar | S.P. Cover, pers. comm. |
| **Subfamily Myrmicinae** |  |  |  |
| ***Veromessor*** |  |  |  |
| *andrei* (Mayr) | 40.0 ± 3.6 (15) | variable | [55, 56] |
| *chamberlini* (Wheeler) | 41.7 ± 5.4 (6) | diurnal | M. Bennett, pers. comm.; R.A. Johnson, pers. obs. |
| *chicoensis* Smith | 44.1 ± 2.1 (6) | variable | [57] |
| *julianus* (Pergande) | 39.0 ± 6.0 (6) | crepuscular-nocturnal-matinal | [58] |
| *lariversi* Smith | 67.8 ± 1.2 (9) | nocturnal | R.A. Johnson, pers. obs. |
| *logobnathus* (Andrews) | 52.1 ± 5.7 (6) | variable | [59, 60]; M. Bennett, pers. comm. |
| *pergandei* (Mayr) | 22.1 ± 2.0 (4) | variable | [53, 60]; R.A. Johnson, pers. obs. |
| RAJ-*pseu* | 69.6 ± 1.6 (8) | nocturnal | [60, as *V. lariversi*]; R.A. Johnson, pers. obs. |
| *smithi* Cole | 51.9 ± 4.3 (6) | crepuscular-nocturnal | [60, 61]; M. Bennett, pers. comm. |
| *stoddardi* (Emery) | 45.1 ± 2.7 (4) | crepuscular-nocturnal | M. Bennett, pers. comm. |

**Table 2.**
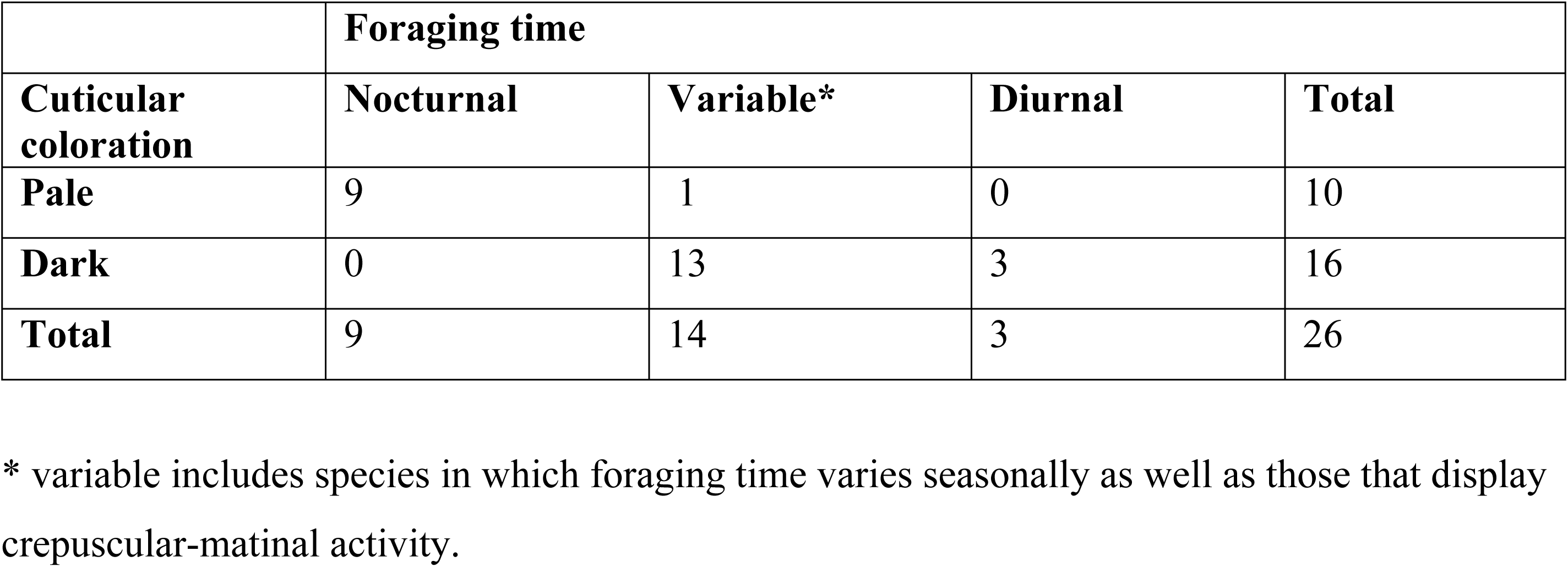
Association between color and activity period based on data in Table 1. Pale species have a brightness (B) value > 65 and dark species have a B value < 60 as measured in Adobe Photoshop (see text).

|  | Foraging time |  |  |  |
| --- | --- | --- | --- | --- |
| Cuticular coloration | Nocturnal | Variable* | Diurnal | Total |
| Pale | 9 | 1 | 0 | 10 |
| Dark | 0 | 13 | 3 | 16 |
| Total | 9 | 14 | 3 | 26 |

\* variable includes species in which foraging time varies seasonally as well as those that display crepuscular-matinal activity.

### Eye area, facet number, and facet diameter

We determined the magnitude of interspecific differences in eye area, facet number, and facet diameter for species in each of the four genera. Before running the MANCOVA for each genus, we tested the homogeneity of variance-covariance (Box’s M test and Levene’s test) and homogeneity of slopes assumptions (Wilks’ lambda for the species × mesosoma interaction effect). Dependent variables met both assumptions for *Myrmecocystus*, *Aphaenogaster*, and *Temnothorax*, but the Levene’s test was significant for *Veromessor* (Table 3). We adjusted for this effect by using a *P* value of 0.01 in our pairwise comparisons for species of *Veromessor*.

**Table 3.** Results for the homogeneity of variance-covariance and regression slopes assumptions in the MANCOVA. The MANCOVA for each genus was run twice; the first run tested homogeneity of variance-covariance and homogeneity of regression slopes assumptions, and the second run was for results of the model. For the Levene’s test, the first column gives the *P* value for the assumptions run, the second gives the *P* value for the results run. Values for these three lines are eye area, facet number, and facet diameter, respectively.

| Genus | Homogeneity of variance-covariance tests |  |  |  | Homogeneity of regression slopes test |  |  |
| --- | --- | --- | --- | --- | --- | --- | --- |
| | Box's M test | $P$ | Levene's test* | | Wilks' $\lambda$ | F | $P$ |
|  |  |  | Assumptions | Results |  |  |  |
| *Myrmecocystus* | $F_{30, 8480} = 1.63$ | 0.016 | 0.047<br>0.048<br>0.13 | 0.10<br>0.38<br>0.14 | 0.698 | $F_{15, 166} = 1.54$ | 0.095 |
| *Aphaenogaster* | $F_{12, 5277} = 0.98$ | 0.47 | 0.20<br>0.38<br>0.28 | 0.12<br>0.21<br>0.89 | 0.743 | $F_{9, 68} = 0.99$ | 0.46 |
| *Temnothorax* | $F_{12, 728} = 2.60$ | 0.002 | 0.62<br>0.27<br>0.55 | 0.18<br>0.60<br>0.55 | 0.522 | $F_{6, 42} = 2.68$ | 0.027 |
| *Veromessor* | $F_{54, 18615} = 1.83$ | 0.0002 | 0.016<br>< 0.001<br>0.44 | 0.16<br>0.012<br>0.48 | 0.775 | $F_{27, 325} = 1.10$ | 0.34 |

#### Myrmecocystus

Eye structure (eye area, facet number, facet diameter) varied significantly across species of *Myrmecocystus* (Wilks’ λ = 0.028, F_15,180_ = 31.9, *P* < 0.001). The tests of between-subject effects demonstrated that all three dependent variables varied significantly across species (eye area: F_5.67_ = 81.5, *P* < 0.001; facet number: F_5,67_ = 4.9, *P* < 0.001; mean *D*: F_5,67_ = 130.0, *P* < 0.001; Fig 6). Based on the estimated marginal means, pairwise comparisons across all species pairs using a LSD test showed that eye area and mean *D* were significantly larger in the pale species (*M. christineae*, *M. navajo*, *M. mexicanus*-02) than in their dark congeners (***M. yuma***, ***M. kennedyi***, ***M. mendax*-03**) (*P* < 0.05, Fig 6). Facet number varied across species in a different manner being significantly higher in *M. mexicanus*-02 and ***M*. *yuma***, but with ***M. yuma*** also overlapping the four other congeners with fewer facets. Relative to the three size-paired pale and dark species (*M. christineae* vs. ***M. yuma***; *M. navajo* vs. ***M. kennedyi***; *M*. *mexicanus*-02 vs ***M. mendax*-03**), mean *D* (using estimated marginal means) was 1.44× larger for *M. christineae* (20.81 μm) compared to ***M. yuma*** (14.42 μm), 1.43× larger for *M. navajo* (20.41 μm) compared to ***M. kennedyi*** (14.29 μm), and 1.41× larger for *M. mexicanus*-02 (20.78 μm) compared to ***M. mendax*-03** (14.79 μm) (Table 4).

**Fig. 6.**
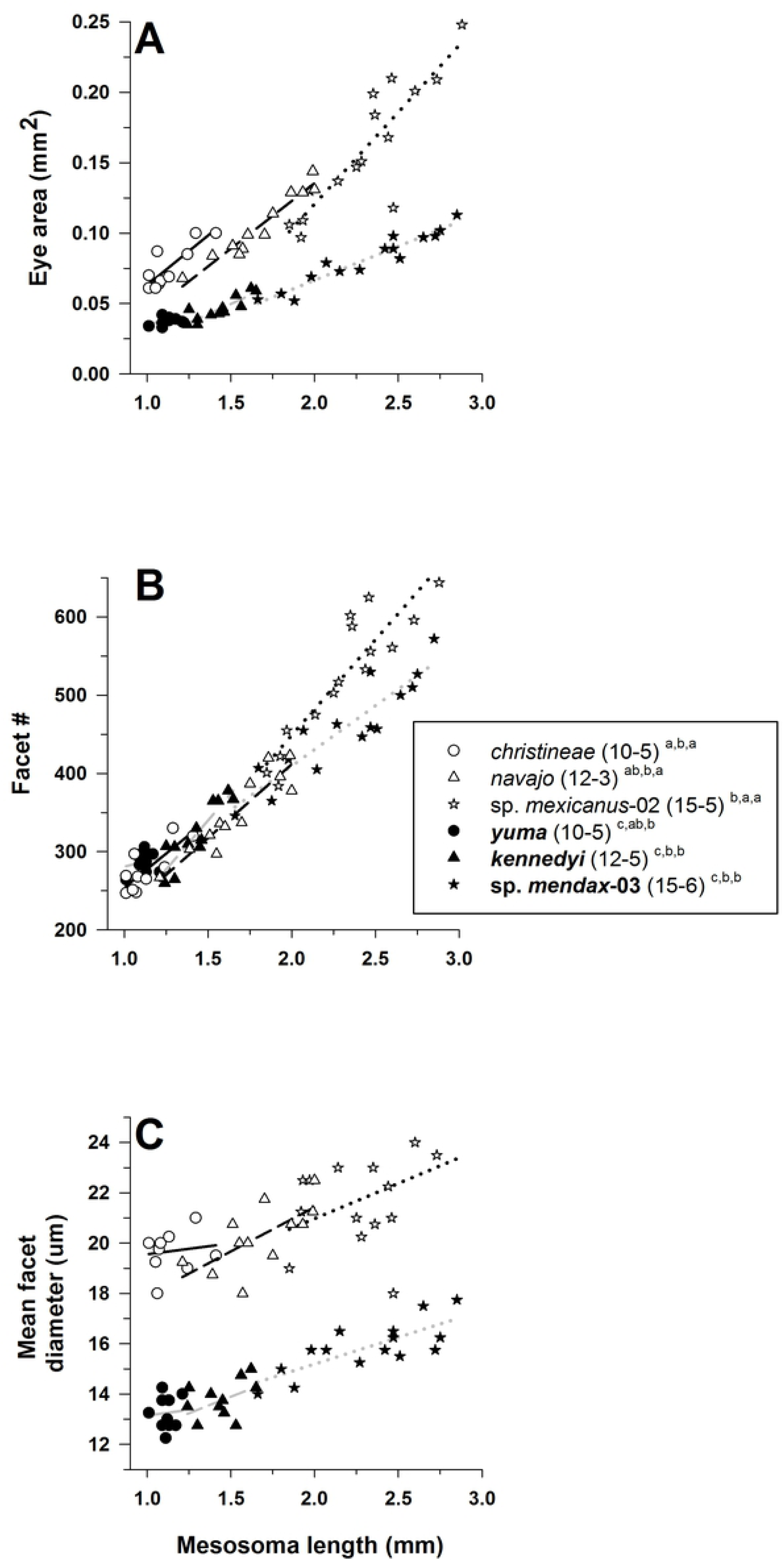
Eye area (mm^2^) (A), facet number (B), and mean facet diameter (D) (µm) (C) for species of *Myrmecocyl·tus* (subfamily Formicinae: tribe Lasiini). Threespecies are pale (open symbols and normal font: *M christineae, M navajo, M mexicanus-*02), and three species aredark (filled symbols and **bold** font: *M yuma, M kennedy,iM mendax-*03) (see text). For each species, number of workers examined and number of colonies they were derived from is given in parentheses. Significant differences *(P* < 0.05) among species are denoted after each species name by the letters *a* c: *a > b* > *c;* the three sets of letters for each species correspond to panels A, B, and C, respectively. Groupings are based on univariate F tests within MANCOVA using the estimated marginal means followed by pairwise comparisons using a least significant differences test (see text). Foraging time for each species is given in Table I.

**Table 4.** Mean values (± 1 SE) for eye area (mm2), number of facets, facet diameter (μm), and mesosoma length (mm) for species examined in this study (see also Figs 6 & 8–10). For each species, values on the first line are raw data, values on second line are estimated marginal mean values using mesosoma as a covariate. Pale species in normal font; dark species in **bold** font (see text).

| Species | Eye area | Number of facets | Facet diameter | Mesosoma length |
| --- | --- | --- | --- | --- |
| *Myrmecocystus*<sup>a</sup> |  |  |  |  |
| *christineae* | 0.0763 ± 0.0049<br>0.1196 ± 0.0077 | 277.5 ± 9.3<br>384.1 ± 14.2 | 19.68 ± 0.26<br>20.81 ± 0.49 | 1.14 ± 0.04 |
| *navajo* | 0.1052 ± 0.0068<br>0.1103 ± 0.0052 | 349.8 ± 14.5<br>352.3 ± 9.6 | 20.27 ± 0.37<br>20.41 ± 0.33 | 1.67 ± 0.07 |
| sp. *mexicanus-02* | 0.1609 ± 0.0118<br>0.1207 ± 0.0067 | 524.1 ± 21.3<br>425.1 ± 12.4 | 21.83 ± 0.50<br>20.78 ± 0.43 | 2.31 ± 0.08 |
| ***yuma*** | 0.0378 ± 0.0009<br>0.0825 ± 0.0079 | 287.0 ± 4.2<br>397.1 ± 14.4 | 13.25 ± 0.21<br>14.42 ± 0.50 | 1.12 ± 0.02 |
| ***kennedyi*** | 0.0463 ± 0.0025<br>0.0685 ± 0.0058 | 322.8 ± 11.3<br>377.6 ± 10.7 | 13.71 ± 0.22<br>14.29 ± 0.37 | 1.43 ± 0.04 |
| **sp. *mendax-03*** | 0.0817 ± 0.0048<br>0.0413 ± 0.0067 | 457.4 ± 16.3<br>358.2 ± 12.4 | 15.85 ± 0.26<br>14.79 ± 0.43 | 2.31 ± 0.10 |
| *Aphaenogaster*<sup>b</sup> |  |  |  |  |
| *megommata* | 0.0815 ± 0.0032<br>0.0745 ± 0.0020 | 241.4 ± 7.9<br>226.1 ± 5.7 | 23.10 ± 0.23<br>22.78 ± 0.31 | 1.68 ± 0.03 |
| *occidentalis* | 0.0264 ± 0.0008<br>0.0328 ± 0.0019 | 79.8 ± 2.1<br>93.7 ± 5.4 | 19.58 ± 0.26<br>19.87 ± 0.30 | 1.42 ± 0.03 |
| *patruelis* | 0.0331 ± 0.0019<br>0.0388 ± 0.0019 | 104.1 ± 4.9<br>116.6 ± 5.3 | 19.96 ± 0.26<br>20.22 ± 0.29 | 1.44 ± 0.02 |
| *boulderensis* | 0.0480 ± 0.0000<br>0.0176 ± 0.0067 | 147.5 ± 3.5<br>81.3 ± 18.7 | 19.88 ± 0.38<br>18.49 ± 1.03 | 2.14 ± 0.02 |
| *Temnothorax*<sup>c</sup> |  |  |  |  |
| sp. BCA-5 | 0.0238 ± 0.0026<br>0.0201 ± 0.0011 | 92.2 ± 6.9<br>80.8 ± 3.7 | 18.30 ± 0.29<br>17.76 ± 0.39 | 0.93 ± 0.06 |
| *neomexicanus* | 0.0109 ± 0.0002<br>0.0128 ± 0.0006 | 66.3 ± 0.9<br>72.0 ± 2.1 | 13.17 ± 0.16<br>14.43 ± 0.23 | 0.67 ± 0.02 |
| *tricarinatus* | 0.0138 ± 0.0003<br>0.0135 ± 0.0005 | 89.0 ± 2.1<br>88.1 ± 1.7 | 13.46 ± 0.23<br>13.42 ± 0.10 | 0.77 ± 0.02 |
| *Veromessor*<sup>d</sup> |  |  |  |  |
| RAJ-pseu | 0.0910 ± 0.0016<br>0.1112 ± 0.0022 | 250.6 ± 3.6<br>283.7 ± 4.9 | 21.17 ± 0.26<br>22.04 ± 0.29 | 1.47 ± 0.18 |
| *lariversi* | 0.0796 ± 0.0027<br>0.0920 ± 0.0019 | 223.5 ± 4.9<br>243.7 ± 4.4 | 21.00 ± 0.31<br>21.53 ± 0.26 | 1.61 ± 0.05 |
| *smithi* | 0.1010 ± 0.0035<br>0.1032 ± 0.0020 | 240.1 ± 5.8<br>243.6 ± 4.5 | 22.52 ± 0.32<br>22.61 ± 0.26 | 1.79 ± 0.04 |
| *pergandei* | 0.0805 ± 0.0049<br>0.0771 ± 0.0018 | 231.9 ± 8.8<br>226.3 ± 4.0 | 19.87 ± 0.26<br>19.72 ± 0.24 | 1.90 ± 0.06 |
| *lobognathus* | 0.0808 ± 0.0024<br>0.0752 ± 0.0020 | 223.8 ± 2.4<br>214.7 ± 4.5 | 20.98 ± 0.33<br>20.74 ± 0.27 | 1.94 ± 0.05 |
| *julianus* | 0.0771 ± 0.0038<br>0.0629 ± 0.0020 | 198.8 ± 7.7<br>175.5 ± 4.5 | 22.07 ± 0.27<br>21.46 ± 0.26 | 2.09 ± 0.06 |
| *chicoensis* | 0.0678 ± 0.0062<br>0.0629 ± 0.0019 | 167.6 ± 10.1<br>159.6 ± 4.2 | 21.00 ± 0.26<br>20.79 ± 0.25 | 1.92 ± 0.09 |
| *andrei* | 0.0663 ± 0.0045<br>0.0492 ± 0.0022 | 198.4 ± 9.1<br>170.5 ± 4.9 | 21.00 ± 0.29<br>20.27 ± 0.29 | 2.14 ± 0.06 |
| *chamberlini* | 0.0395 ± 0.0013<br>0.0513 ± 0.0021 | 126.7 ± 3.2<br>145.9 ± 4.7 | 19.60 ± 0.23<br>20.11 ± 0.28 | 1.62 ± 0.03 |
| *stoddardi* | 0.0483 ± 0.0035<br>0.0476 ± 0.0018 | 130.7 ± 6.4<br>129.5 ± 4.0 | 20.43 ± 0.31<br>20.40 ± 0.24 | 1.85 ± 0.08 |

<sup>a</sup> = estimated marginal means evaluated at a mesosoma length of 1.7434 mm.
<sup>b</sup> = estimated marginal means evaluated at a mesosoma length of 1.5468 mm.
<sup>c</sup> = estimated marginal means evaluated at a mesosoma length of 0.7534 mm.
<sup>d</sup> = estimated marginal means evaluated at a mesosoma length of 1.8333 mm.

Mesosoma length also was a significant covariate in the model (Wilks’ λ = 0.400, F_3, 65_ = 32.5, *P* < 0.001), and tests of between-subjects effects were significant for all three variables (eye area: F_1,67_ = 85.5, *P* < 0.001; facet number: F_1,67 df_ = 95.3, *P* < 0.001; mean *D*: (F_1,67_ = 12.9, *P* < 0.001). All three eye features increased with body size within all six species (Fig 6).

The ANCOVA for diameter of the anterior ocellus was significant (Fig 7; tests of between-subject effects: F_7,98_ = 69.6, *P* < 0.001), but the species × mesosoma length interaction term was not significant (F_7,91_ = 0.96, *P* > 0.45). Pairwise comparisons between all species pairs using LSD tests showed that anterior ocellus diameter usually was highest for dark species, though the diameter for one pale species (*M. christineae*) overlapped with this group. The other four pale species (*M*. *testaceus*, *M. mexicanus*-01, *M. mexicanus*-02, *M. navajo*) were significantly lower than all other congeners (Fig 7). Note that *M. navajo* was not included in our statistical analysis, but it was placed lowest in this group post-hoc because the anterior ocellus was lacking in 11 of 12 workers.

**Fig. 7.**
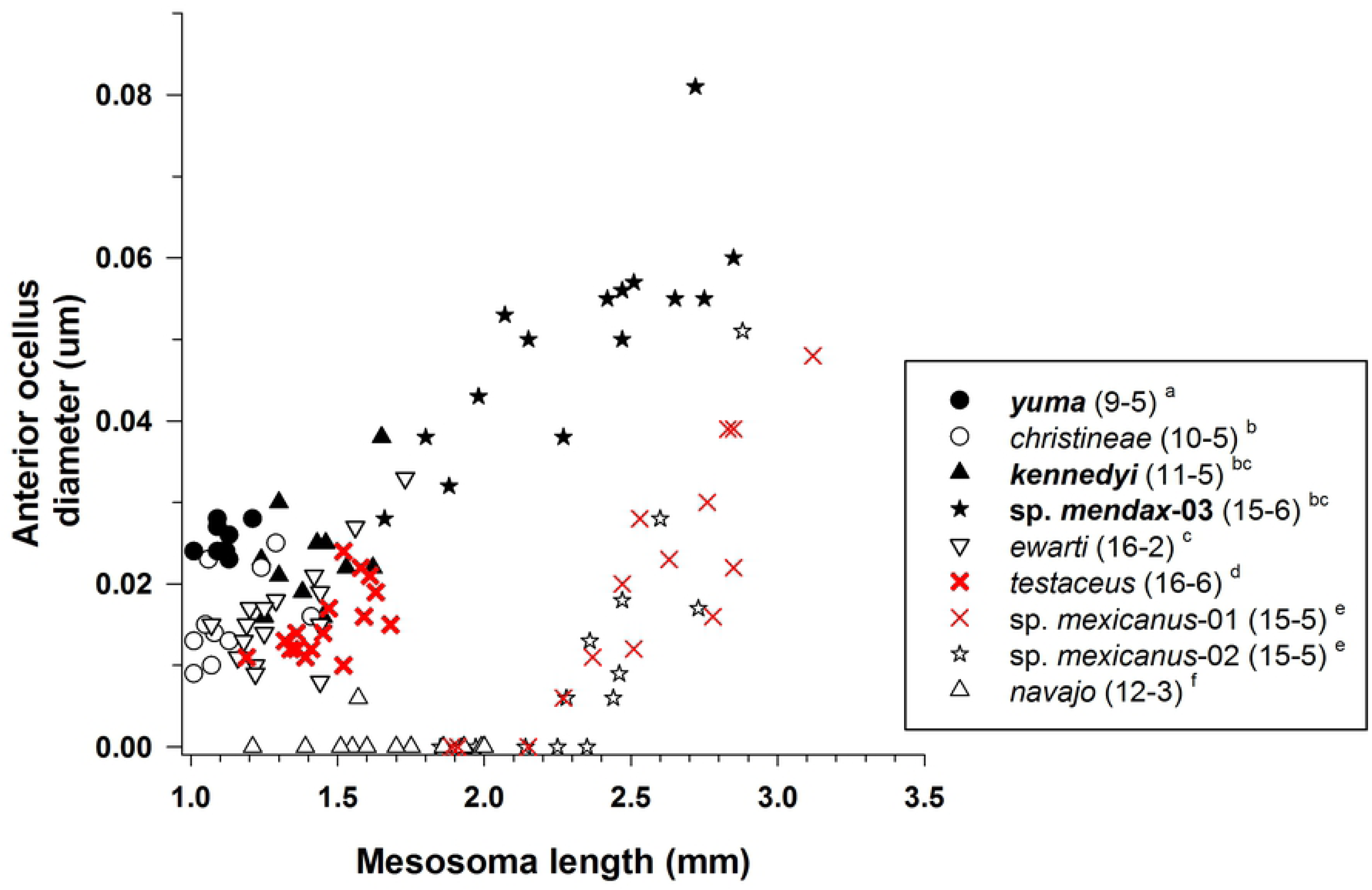
Anterior ocellus diameter for species of *Myrmecocy.ftu s* ( s ubfamily Formicinae: tribe Lasiini). Six species are pale (open or red symbols and normal font: *M chrisLineae, M ewarli, M navajo, M. testaceu s, M m e xicanu s-0* l, *M. mexicanu s-0 2 ),* and three species are dark (filled symbols and **bold** font: *M. yuma, M. kennedyi, M. mendax-03 )* (see text). For each species, number of workers examined and number of colonies they were derived from is given in parentheses. Significan t differences *(P* < 0.05) among species are denoted after each species name by the letters *a-f a > b* > *c* > *d* > *e* > *f,* the three sets of letters for each species correspond to panels A, B, and C, respectively. Groupings are based on univariate F tests within ANCOVA followed by pairwise comparisons of the estimated marginal means using a least significant differences test (see text). Foraging time for each species is given in Table I.

Diameter of the anterior ocellus also increased with body size within all species except *M. navajo*. Presence of the anterior ocellus also was associated with body size in *M. mexicanus*- 01 and *M. mexicanus*-02, as workers with a mesosoma length < ≈ 2.2 mm lacked an anterior ocellus while those with a mesosoma length > ≈ 2.2 mm had this ocellus, with ocellus diameter increasing with body size in these latter workers (Fig 7). Both posterior ocelli usually were present, but tiny, in workers that lacked an anterior ocellus.

#### Aphaenogaster

Eye structure (eye area, facet number, facet diameter) varied significantly across species of *Aphaenogaster* (MANCOVA: Wilks’ λ= 0.042, F_9,76_ = 22.5, *P* < 0.001). Tests of between-subject effects demonstrated that all three dependent variables varied significantly across species (eye area: F_3,33_ = 125.0, *P* < 0.001; facet number: F_3,33_ = 146.8, *P* < 0.001; mean *D*: F_3,33_ = 31.9, *P* < 0.001; Fig 8). Based on the estimated marginal means, pairwise comparisons across all species pairs using an LSD test showed that all three eye measures were significantly higher for the pale *A. megommata* than for all three dark congeners (*P* < 0.01). Mean *D* (using estimated marginal means) was 1.15× larger for *A. megommata* (22.78 μm) compared to ***A. occidentalis*** (19.87 μm), 1.13× larger than that for ***A. patruelis*** (20.22 μm), and 1.23× larger than that for ***A. boulderensis*** (18.49 μm) (Table 4).

**Fig. 8.**
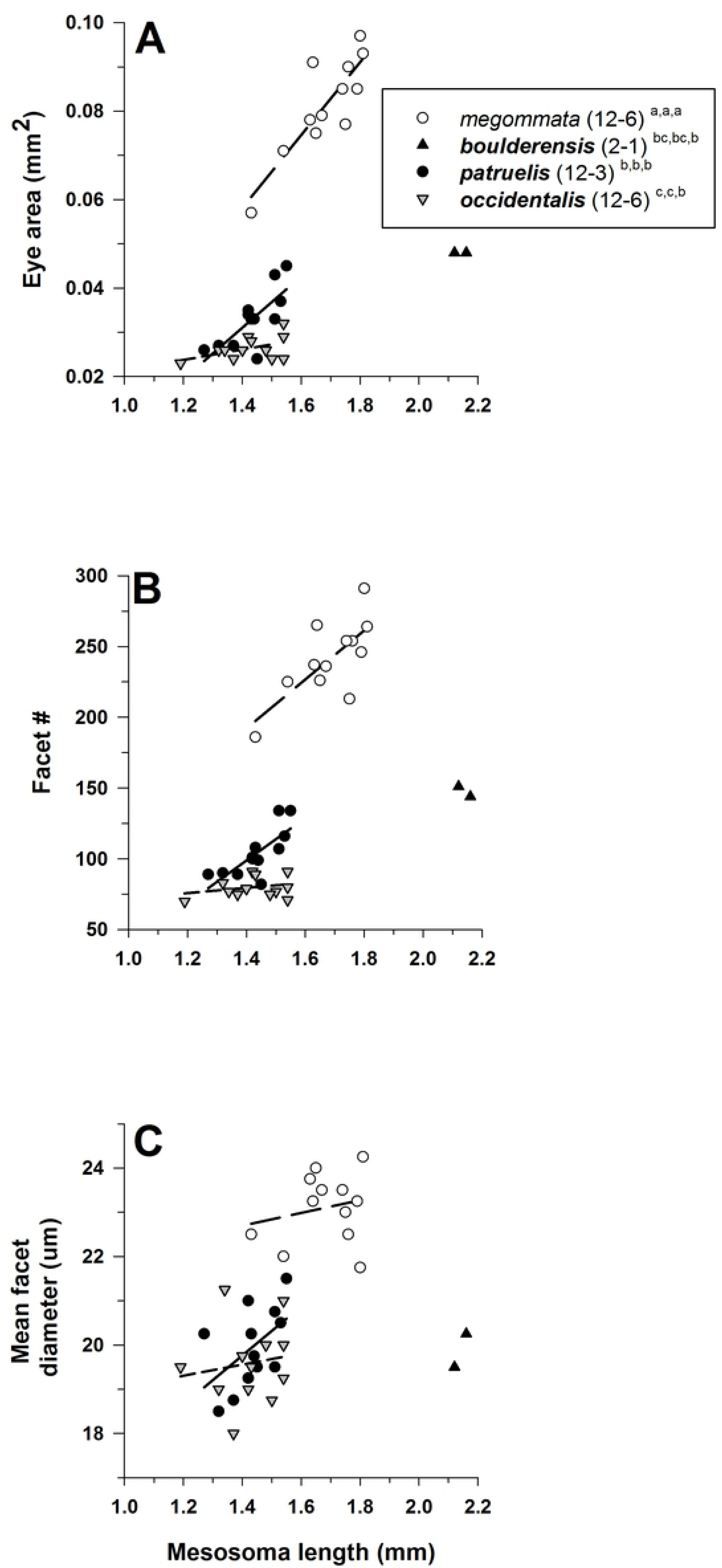
Eye area (mm^2^) (A), facet number (B), and mean facet diameter (D) (µm) (C) for species of *Aphaenogaster* (subfamily Myrmicinae: tribe Stenammini). *Aphaenogaster megommata* is pale (open symbols and regular font), while *A. boulderensis, A. occidentalis,* and *A. patruelis* are dark (filled symbols and **bold** font) (see text). For each species, number of workers examined and number of colonies they were derived from is given in parentheses. Significant differences *(P* < 0.05) among species aredenoted after each species name by the letters *a-c : a > b* > *c;* the three sets of letters for each species correspond to panels A, B, and C, respectively. Groupings are based on univariate F tests within MANCOVA using the estimated marginal means followed by pairwise comparisons using a least significant differences test (see text). Foraging time for each species is given in Table I.

Mesosoma length was a significant covariate in the model (Wilks’ λ = 0.502, F_3,31_ = 10.2, *P* < 0.001), and tests of between-subjects effects were significant for eye area (F_1,33_ = 28.1, *P* < 0.001) and facet number (F_1,33_ = 17.1, *P* < 0.001), but not for mean *D* (F_1,33_ = 2.7, *P* = 0.11). These patterns were evidenced in that eye area and facet number increased with body size within all three species (Fig 8; ***A. boulderensis*** excluded because of small sample size), while mean *D* increased with body size for *A. megommata* and ***A. patruelis***, but it decreased with body size for ***A*. *occidentalis*** (Fig 8).

#### Temnothorax

Eye structure (eye area, facet number, facet diameter) varied significantly across species of *Temnothorax* (MANCOVA: Wilks’ λ= 0.039, F_6,46_ = 31.0, *P* < 0.001). The tests of between-subject effects demonstrated that all three dependent variables varied significantly across species (eye area: F_2,25_ = 24.9, *P* < 0.001; facet number: F_2,25_ = 20.8, *P* < 0.001; mean *D*: F_2,25_ = 53.4, *P* < 0.001; Fig 9). Based on the estimated marginal means, pairwise comparisons across all species pairs using a LSD test showed that eye area and *D* were significantly larger for the pale *T.* sp. BCA-5 than for the two dark congeners. Facet number was highest in ***T*. *tricarinatus***, lowest in ***T. neomexicanus***, and intermediate to and overlapping both other species for *T*. sp. BCA-5 (Fig 9). Mean *D* (using estimated marginal means) was 1.23× larger for *T.* sp. BCA-5 (17.76 μm) compared to ***T. neomexicanus*** (14.43 μm) and 1.32× larger than that for ***T. tricarinatus*** (13.42 μm) (Table 4).

**Fig. 9.**
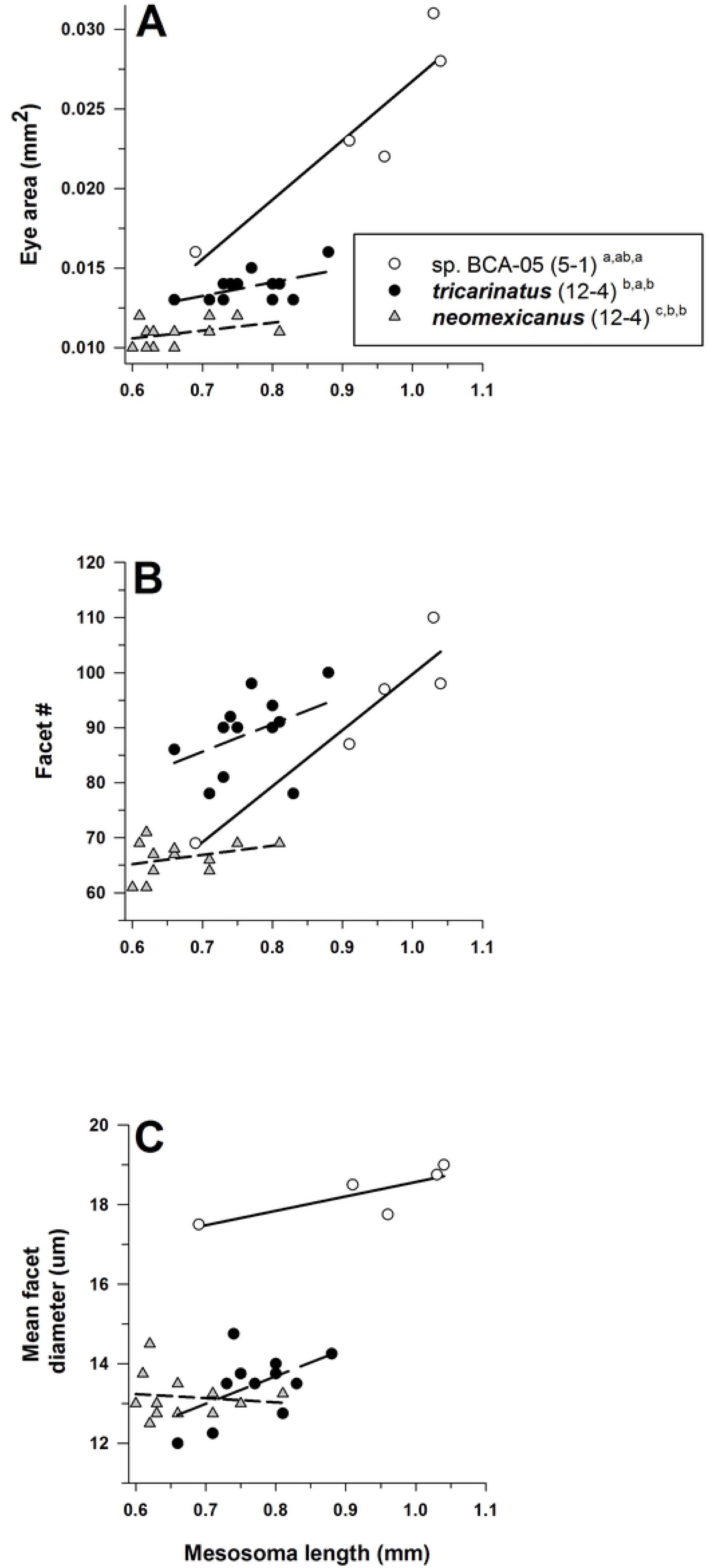
Eye area (mm^2^) (A), facet number (B), and mean facet diameter (D) (µm) (C) for species of *Temnothorax* (subfamily Myrmicinae: tribe Crematogastrini). *Temnothorax* sp. BCA-S is pale (open symbols and regular font), wh.ile *T. neomexicanus* and *T. triacarinatus* aredark (filled symbols and **bold** font) (see text). For each species, number of workers examined and number of colonies they were derived from is given in parentheses. Significant differences *(P* < 0.05) among species are denoted after each species name by the letters *a-b: a> b;* the three setsof letters for each species correspond to panels A, B, and C, respectively. Groupings are based on univariate F tests within MANCOVA using the estimated marginal means followed by pairwise comparisons using a least significant differences test (see text). Foraging time for each species is given in Table I.

Mesosoma length was a significant covariate in the model (Wilks’ λ = 0.448, F_3,23_ = 9.45, *P* < 0.001), and tests of between-subjects effects were significant for eye area (F_1,25_ = 29.9, *P* < 0.001) and facet number (F_1,25_ = 20.7, *P* < 0.001), but not for mean *D* (F_1,25_ = 3.9, *P* = 0.058). Eye area and facet number increased with body size within all three species, while mean *D* increased with body size for *T*. sp. BCA-5 and ***T. tricarinatus***, but it decreased with body size for ***T. neomexicanus*** (Fig 9).

#### Veromessor

Eye structure (eye area, facet number, facet diameter) varied significantly across species of *Veromessor* (MANCOVA: Wilks’ λ= 0.020, F_27,351_ = 36.5, *P* < 0.001). The tests of between-subject effects demonstrated that all three dependent variables varied significantly across species (eye area: F_9,122_ = 149.2, *P* < 0.001; facet number: F_9,122_ = 141.4, *P* < 0.001; mean *D*: F_9,122_ = 12.5, *P* < 0.001; Fig 10). Based on the estimated marginal means, pairwise comparisons across all species pairs using a LSD test showed that eye area was significantly larger for the pale *V*. RAJ-*pseu* than for the dark ***V. smithi***, and eyes for both species were significantly larger than the pale *V. lariversi*. Eye area was significantly lower for all other dark congeners (Fig 10). Facet number was significantly higher for *V.* RAJ-*pseu* than for ***V. smithi*** and *V. lariversi*, and these three species were all significantly higher than all other dark congeners. Mean *D* (using estimated marginal means) was highest for ***V. smithi*** and *V.* RAJ-*pseu*, followed by *V. lariversi* and ***V. julianus***, with the two latter species overlapping with *V.* RAJ-*pseu* but not ***V. smithi***. Mean *D* was significantly lower for all other dark congeners (*P* < 0.05, Fig 10; Table 4).

**Fig. 10.**
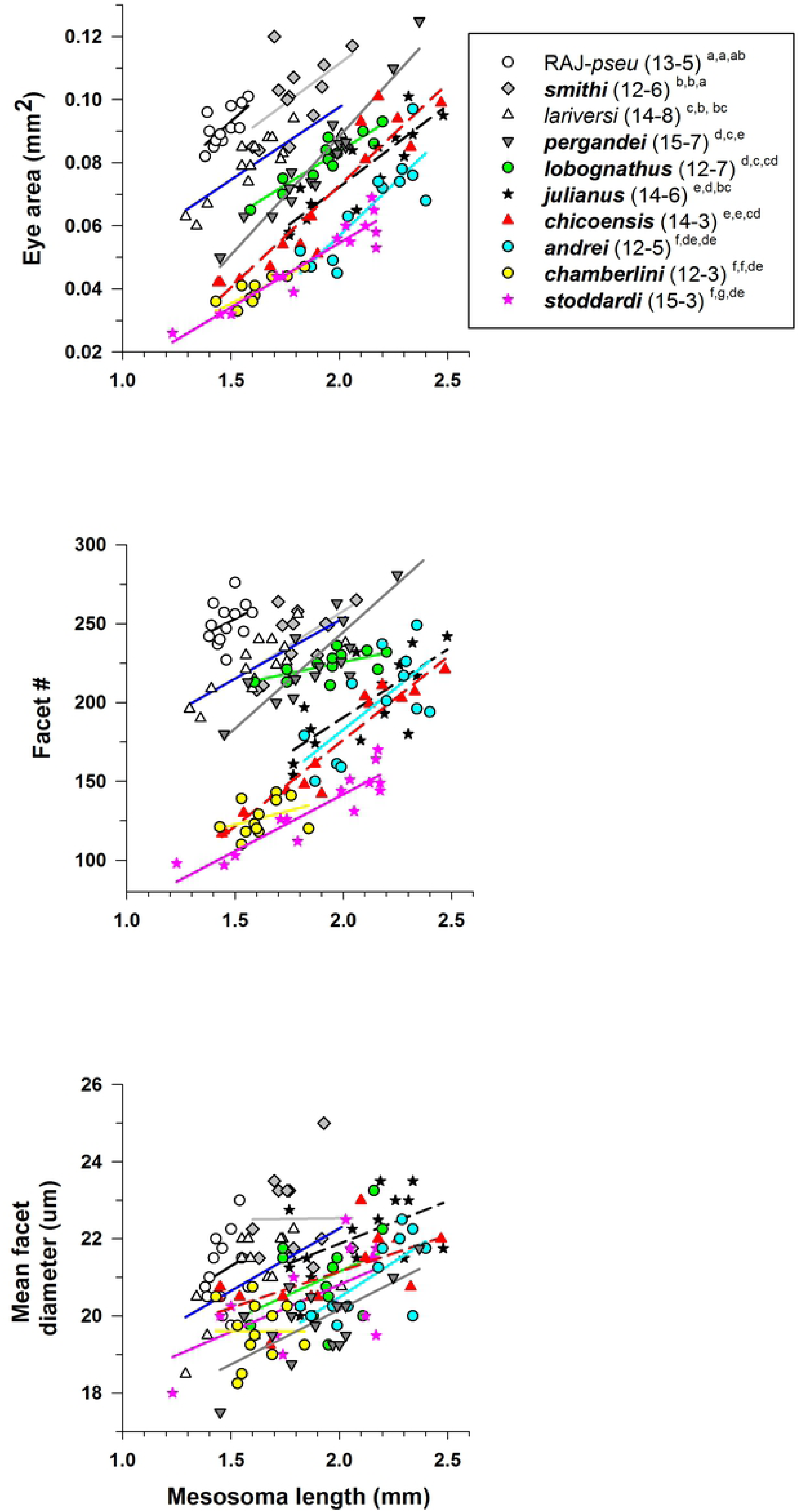
Eye area (mm^2^) (A), facet number (B), and mean facet diameter (D) (µm) (C) for species of *Veromessor* (subfamily Myrmicinae: tribe Stenammini). *Veromessor lariversi* and *V. RAJ-pseu* are pale (open symbols and regular font), while the other eight species are dark (filled symbols and **bold** font)(see text). For each species, number of workers examined and number of colonies they derived from is given in parentheses. Significant differences *(P* < 0.01) among species are denoted after each species name by the letters *a- g: a > b* > *c* > *d* > *e* > *f* > *g;* the three sets of letters for each species correspond to panels A, B, and C, respectively. Groupings are based on univariate F tests within MANCOVA using the estimated marginal means followed by pairwise comparisons using a least significant differences test (see text). Foraging time for each species is given in Table I.

Mesosoma length was a significant covariate in the model (Wilks’ λ = 0.198, F_3,120_ = 162.4, *P* < 0.001), and tests of between-subjects effects were significant for all three variables (eye area: F_1,122_ = 486.0, *P* < 0.001; facet number: F_1,122_ = 149.2, *P* < 0.001; mean *D*: F_1,122_ = 40.9, *P* < 0.001). Eye area and facet number increased with body size within all 10 species of *Veromessor*, and mean *D* increased for all species except ***V. smithi*** and ***V*. *chamberlini*** (Fig 10).

### Detailed Eye Measurements

#### Variation in interommatidial angle (**Δ*ϕ***)

Values of Δ*ϕ* ranged from 3.5–7° among the workers studied (Fig 11). The ANCOVA for Δ*ϕ* was significant for genus (F_3,32_ = 10.1, *P* < 0.001), but not for activity period (F_1,32_ = 4.0, *P* = 0.055); the interaction of genus × activity period was also significant (F_3,32_ = 7.3, *P* = 0.001). The dark species had marginally larger Δ*ϕ*‘s than the pale species (*P* = 0.055), and the significant interaction between Δ*ϕ* and genus indicated significant differences among genera in the direction and magnitude of differences in Δ*ϕ*. Pale species had larger mean Δ*ϕ*’s in *Myrmecocystus* (t-test: t _8_ = -3.4, *P* < 0.02) and *Veromessor* (t = -0.3, *P* > 0.7), but dark species had larger Δ*ϕ*’s in *Aphaenogaster* (t = -3.6, *P* < 0.01) and *Temnothorax* (t = 2.7, *P* < 0.03) (Fig 11). Across genera, Δ*ϕ* was lowest for *Myrmecocystus* and *Veromessor* and greatest for *Aphaenogaster* and *Temnothorax* (Tukey’s HSD test, *P* < 0.05) (Fig 11).

**Fig. 11.**
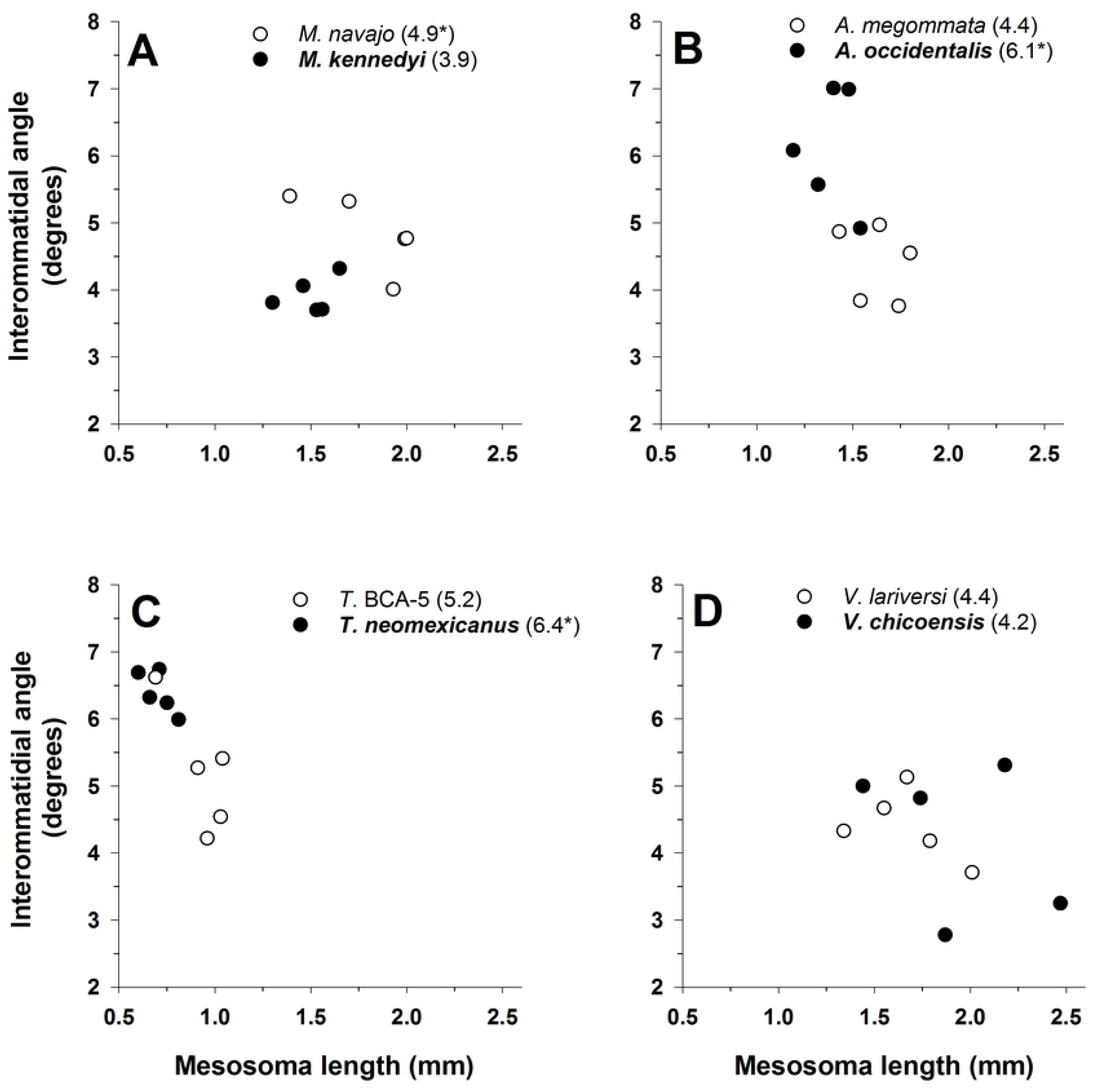
Interommatidial angle (A¢,) for one pale (open circles and regular font) and one dark (filled circles and bold font) (see text) species in each of four ant genera: *(A )Myrmecocystus, (B) Aphaenogaster,* **(C)** *Temnothorax,* and **(D)** *Veromessor.* All plots have the same x-axis and y-axis scaling in order to visualize differences between light and dark species across genera. Mean *A¢* (in degrees) is given after each species name with an asterisk denoting the species with a significant larger *A¢* based on at-test *(P* < 0.05). The significant interaction of genus x activity period is shown by larger */:J.¢ s*for pale species of *Myrmecocystus* and *Veromessor,* whereas */:J.¢* was larger for dark species of *Aphaenogaster* and *Temnothorax.* Sample sizeis *n* = *5* for each species.

We also ran the above model with mesosoma length as a covariate, and it was significant (F_1,31_ = 5.4, *P* = 0.026). This significance largely was caused by Δ*ϕ* decreasing in larger workers of *Temnothorax* and *Veromessor*, while this angle did not vary with mesosoma length within species of *Myrmecocystus* and *Aphaenogaster* (Fig 11).

#### Eye parameter (*ρ*))

The ANCOVA for *ρ* was significant for genus (F_3,32_ = 11.6, *P* < 0.001), activity period (F_1,32_ = 11.2, *P* = 0.002), and the interaction of genus × activity period (F_3,32_ = 13.1, *P* < 0.001). As expected, overall *ρ* was greater for pale (mean = 1.70) than for dark species (mean =1.51), however, a significant genus × activity period interaction indicated differences in direction and magnitude of these differences (Fig 12). Pale species had the larger mean *p* in *Myrmecocystus* (t-test: t_8 df_ = -8.9, *P* < 0.001), *Veromessor* (t_8_ = -0.3, *P* > 0.7), and *Temnothorax* (t_8_ = -1.7, *P* > 0.10), but Δ*ϕ* was larger for the dark species in *Aphaenogaster* (t_8_ = -2.2, *P* < 0.06). Across genera *ρ* was highest for *Aphaenogaster,* intermediate for *Temnothorax* and *Veromessor*, and lowest for *Myrmecocystus* (Tukey’s HSD test, *P* < 0.05; Fig 12).

**Fig. 12.**
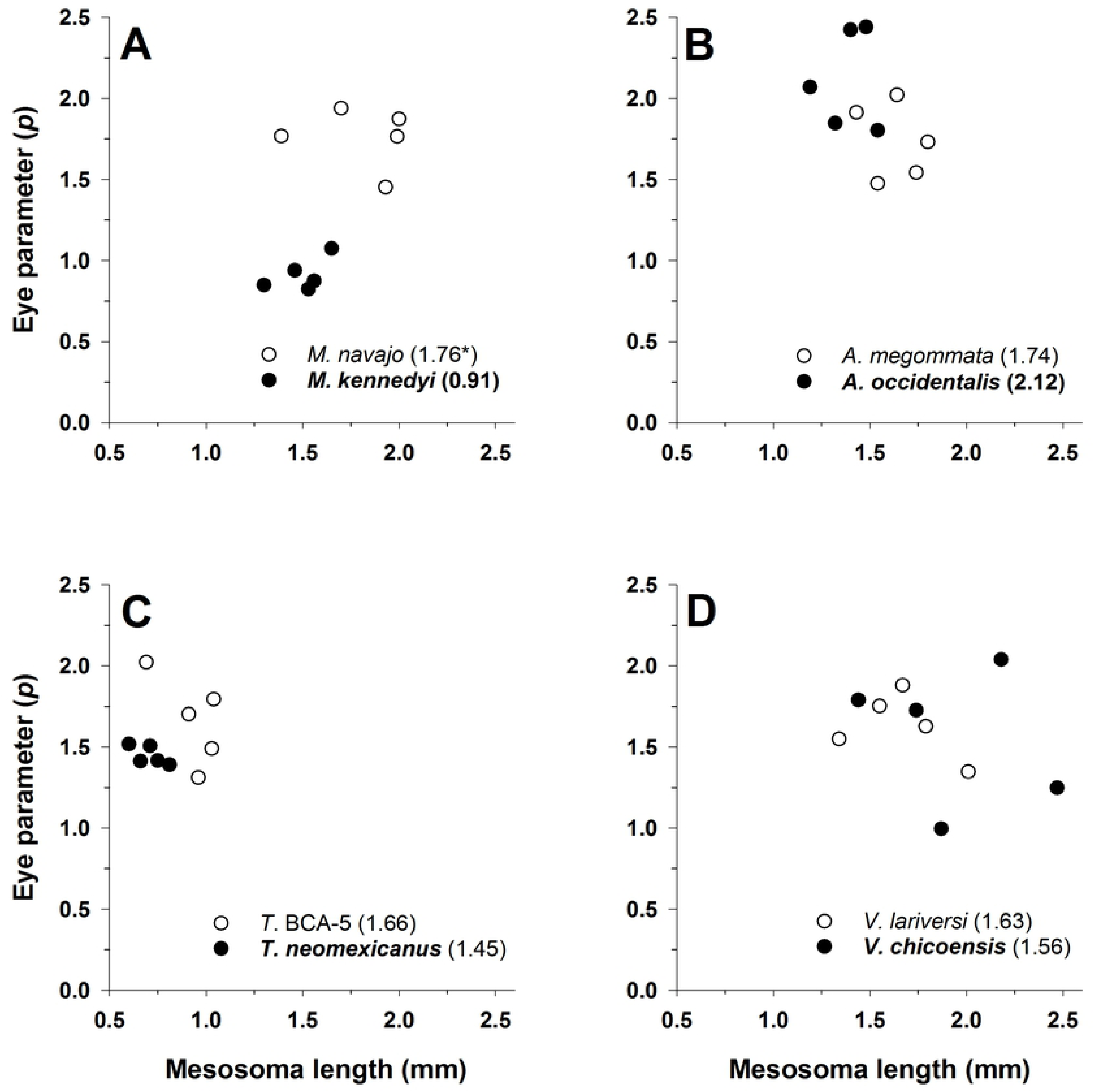
Eye parameter (ρ) for one pale (open circles and regular font) and one dark (filled circles and bold font) (see text) species in each of four ant genera: *(A )Myrmecocystus, (B) Aphaenogaster,* **(C)** *Temnothorax,* and **(D)** *Veromessor.* All plots have the same x-axis and y-axis scaling in order to visualize differences between light and dark species across genera. Mean *p* is given after each species name with an asterisk denoting the species with a significant larger *p* based on at-test *(P* < 0.05). The significant interaction of genus x activity period is shown by larger differences between light-colored and dark-colored species of *Aphaenogaster* compared to those in the other three genera. Sample sizeis *n* = *5* for each species.

We also ran the above model with mesosoma length as a covariate, but it was not significant (F_1,31_ = 1.6, *P* > 0.20). The *ρ* was not positively or negatively correlated with body size for any of the examined species (Fig 11).

#### Visual field span

The ANCOVA for visual field span was significant for genus (F_3,32_ = 53.6, *P* < 0.001), activity period (F_1,32_ = 151.7, *P* < 0.001), and the interaction of genus × activity period (F_3,32_ = 14.8, *P* < 0.001). Visual field span was greater for pale (mean = 98.8°) than for dark species (mean = 73.0°), and the significant genus × activity period interaction indicated that differences between the visual field of pale and dark species were larger in some genera, e.g., *Aphaenogaster*, than others (Fig 13). Though not always significantly different, pale species had a larger mean visual field in all four genera (*Myrmecocystus* t-test t_8 df_ = - 2.90, *P* = 0.10; *Aphaenogaster*: t_8_ = 12.7, *P* < 0.001; *Temnothorax*: t-test t_8_ = -4.6, *P* = 0.002; *Veromessor*: t_8_ = - 6.9, *P* < 0.001). The pale species of *Myrmecocystus* was also significantly different when comparing the means when including mesosoma as a covariate (F_1,7_ = 87.0, *P* < 0.001). Across genera the visual field was greatest for *Myrmecocystus*, intermediate for *Aphaenogaster*, and smallest in *Temnothorax* and *Veromessor* (Tukey’s HSD test, *P* < 0.05; Fig 13).

**Fig. 13.**
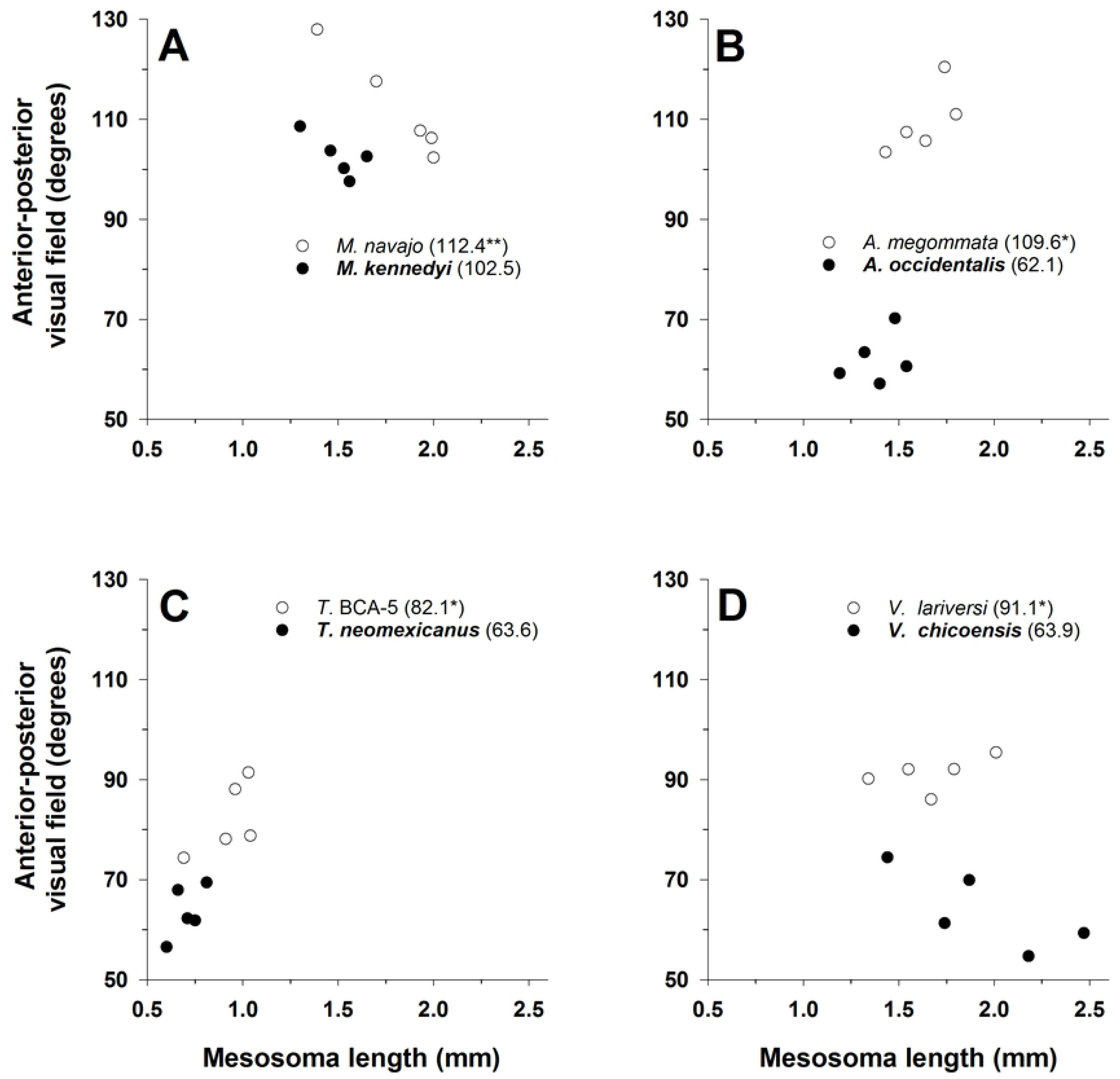
Anterior-posterior visual field span (in degrees) for one pale (open circles and regular font) and one dark (filled circles and bold font) (see text) species in each of four ant genera: **(A)** *Myrmecocystus,* **(B)** *Aphaenogaster,* **(C)** *Temnothorax,* and **(D)** *Veromessor.* All plots have the same x-axis and y-axis scaling in order to visualize differences between pale and dark species across genera. Mean visual field span (in degrees) is given after each species name with an asterisk denoting the species with a significant larger visual field based on at-test *(P* < 0.05); the double asterisk denotes that the t-test was not significant, but that the visual field was significantly larger when including mesosoma length as a covariate. The significant interaction of genus x activity period is shown by larger differences between pale and dark species of *Aphaenogaster* compared to those in the other three genera. Sample sizeis *n* = 5 for each species.

We also ran the above model with mesosoma length as a covariate, but it was not significant (F_1,31_ = 2.6, *P* = 0.11), in part, because of the differing patterns exhibited across species. For example, visual field span was positively correlated with mesosoma length in *A. megommata*, ***A. occidentalis***, *T*. BCA-5, ***T. neomexicanus***, and *V. lariversi*, but these two variables were negatively correlated in ***M. kennedyi***, *M. navajo*, and ***V. chicoensis*** (Fig 13).

The maximum visual span for pale species was 128° for *M. navajo* and 121° for *A. megommata*; the maximum for all other species was < 120°. Because their eyes are located on the side of the head, the center of these relatively small visual field spans was directed laterally. This means that, in the ants examined here, there was no forward part of the visual field for either eye directed toward the mouthparts, and so there was no anterior region of binocular vision.

#### Regional variation in *D*

*D* varied regionally in three (*M. navajo, **V. chicoensis**, V. lariversi*) of the four species; all three species met the assumption of sphericity (Table 5; Fig 14). Based on a post-hoc LSD test, the anterior and ventral facets were largest in *M. navajo*, ventral facets were largest in ***V. chicoensis***, and lateral and ventral facets were largest in *V. lariversi*. *D* did not vary across regions in ***M. kennedyi***, but the ventral facets had the largest mean *D* (Table 5; Fig 14).

**Fig. 14.**
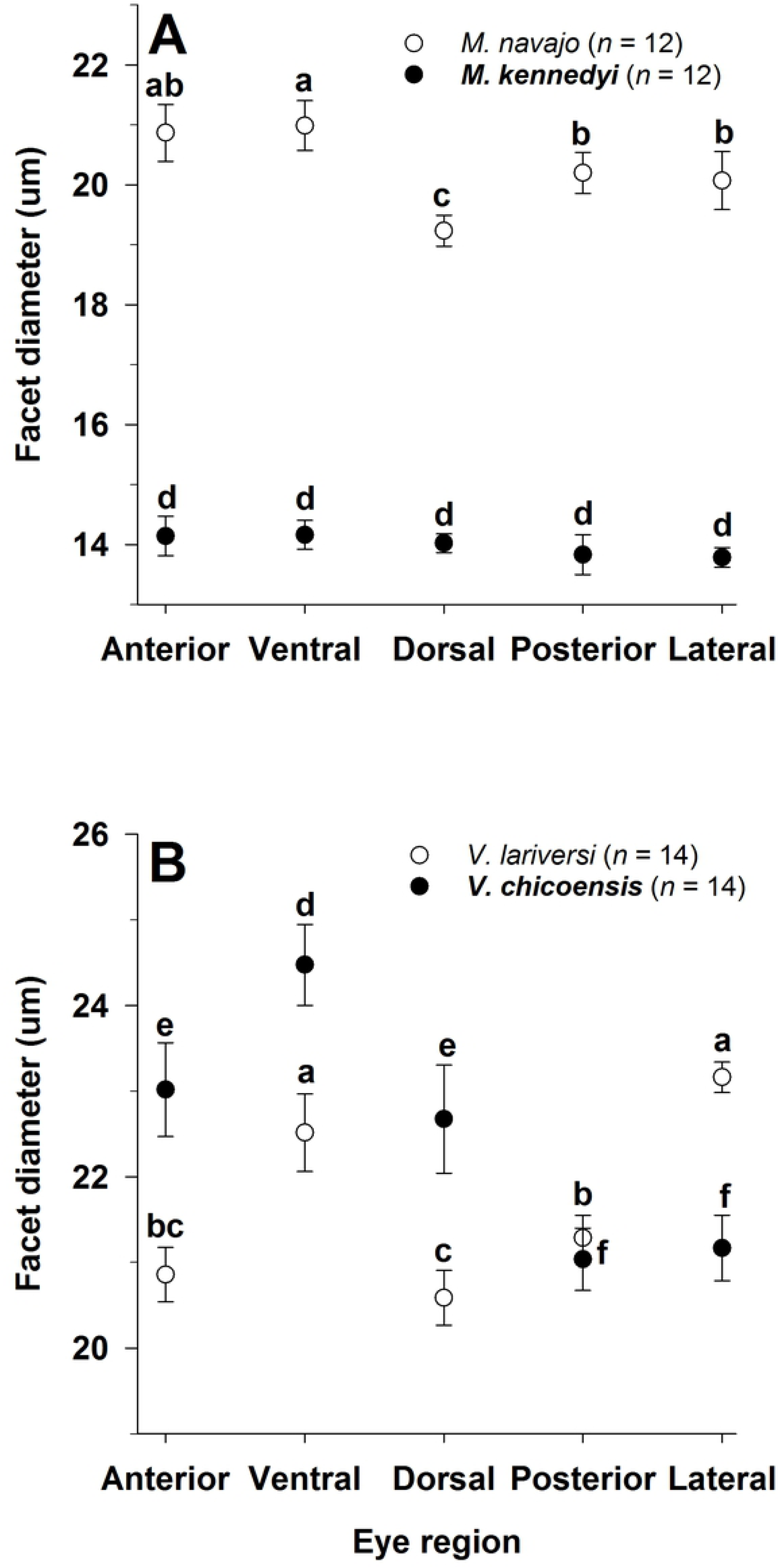
Regional variation in facet diameter for one pale (open circles and regular font) and one dark (filled circles and bold font) (see text) species in each of two ant genera: (A) *Myrmecocystus* and (B) *Veromessor.* Significant differences within each species are denotedby the letters *a-c: a> b* > *c* for pale species; *d-f d* > *e* > *f* for dark species. Groupings for each species are based on a repeated-measures ANOVA followed by a least significant differences test. Sample size is given after each species name.

**Table 5.**
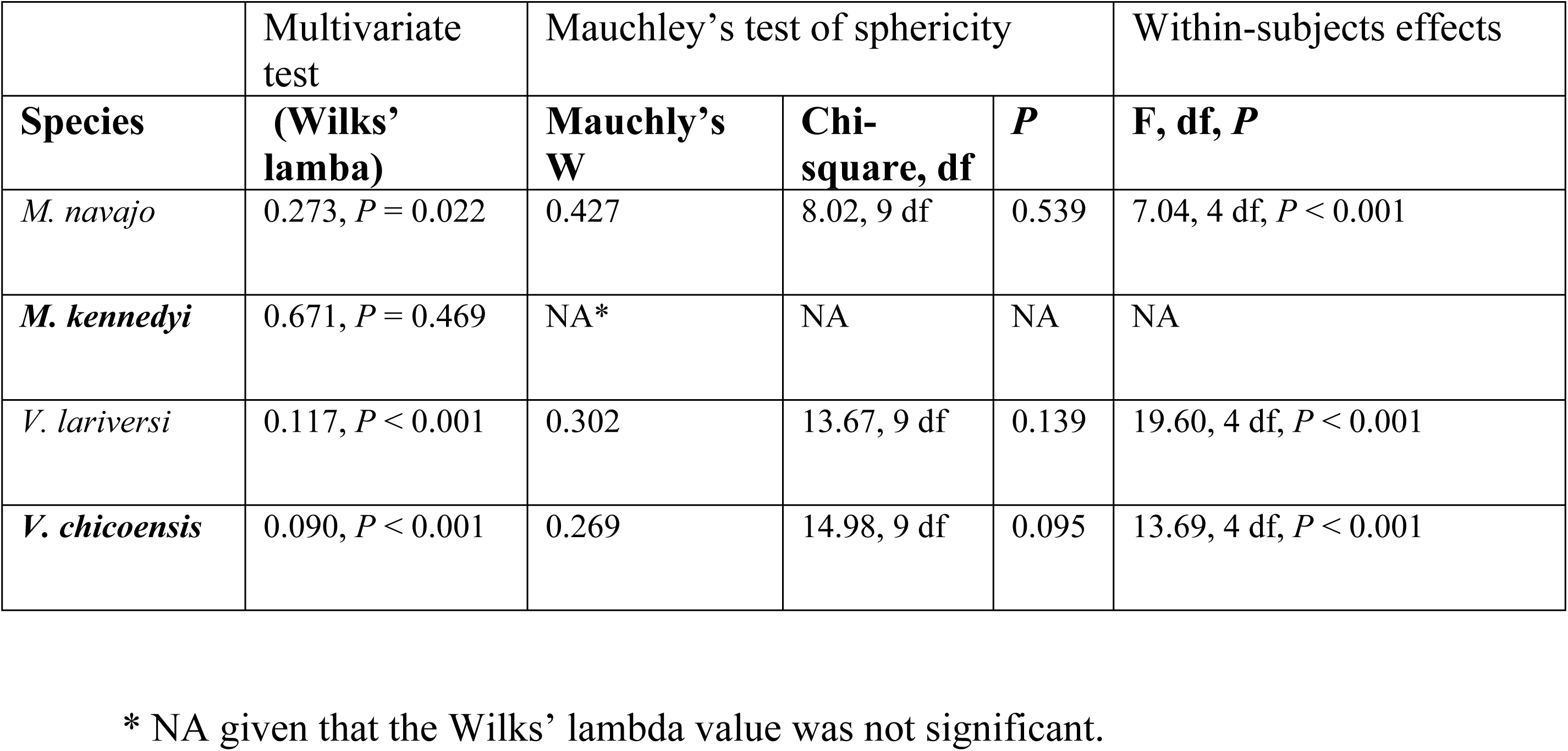
Repeated measures ANOVA results for regional variation in facet diameter for one pale (normal font) and one dark species (**bold** font) (see text) of *Myrmecocystus* and *Veromessor* (*n* = 12 per species for *Myrmecocystus*; *n* = 14 per species for *Veromessor*).

|  | Multivariate test | Mauchly's test of sphericity |  |  | Within-subjects effects |
| --- | --- | --- | --- | --- | --- |
| Species | (Wilks' lamba) | Mauchly's W | Chi-square, df | P | F, df, P |
| *M. navajo* | 0.273, $P = 0.022$ | 0.427 | 8.02, 9 df | 0.539 | 7.04, 4 df, $P < 0.001$ |
| ***M. kennedyi*** | 0.671, $P = 0.469$ | NA* | NA | NA | NA |
| *V. lariversi* | 0.117, $P < 0.001$ | 0.302 | 13.67, 9 df | 0.139 | 19.60, 4 df, $P < 0.001$ |
| ***V. chicoensis*** | 0.090, $P < 0.001$ | 0.269 | 14.98, 9 df | 0.095 | 13.69, 4 df, $P < 0.001$ |

\* NA given that the Wilks' lambda value was not significant.

#### Additional pale ant species with enlarged eyes

We visually identified numerous additional pale ant species with enlarged eyes during our survey on Antweb. We confirmed our visual estimate of brightness by measuring B values on available workers of these species, as detailed above, finding that numerous species displayed a B value greater than 70, which we used as our lower threshold for pale species (Table 6). We also visually judged that eyes for all of these species were larger than that of their dark congeners. The combination of pale color and enlarged eyes occurred in numerous additional species of *Temnothorax* from both the Old and New World, as well as in four additional genera – *Crematogaster* and *Messor* (subfamily Myrmicinae), and *Dorymyrmex* and *Iridomyrmex* (subfamily Dolichoderinae) (Table 6); the latter two genera comprise a third subfamily containing pale species. Moreover, this combination of traits occurred in both the Old and New World, and they were especially common in *Temnothorax* (Table 6), where these traits evolved independently in multiple species groups (in at least two species groups in the United States and Mexico (***T. silvestrii*** and *T. tricarinatus*) and in at least one species group in northern Africa (***T. laciniatus***) (Table 6). Interestingly, several pale species of *Temnothorax* appeared to not have enlarged eyes, e.g., *T. agavicola*, *T. atomus*, and *T. indra*.

**Table 6.**
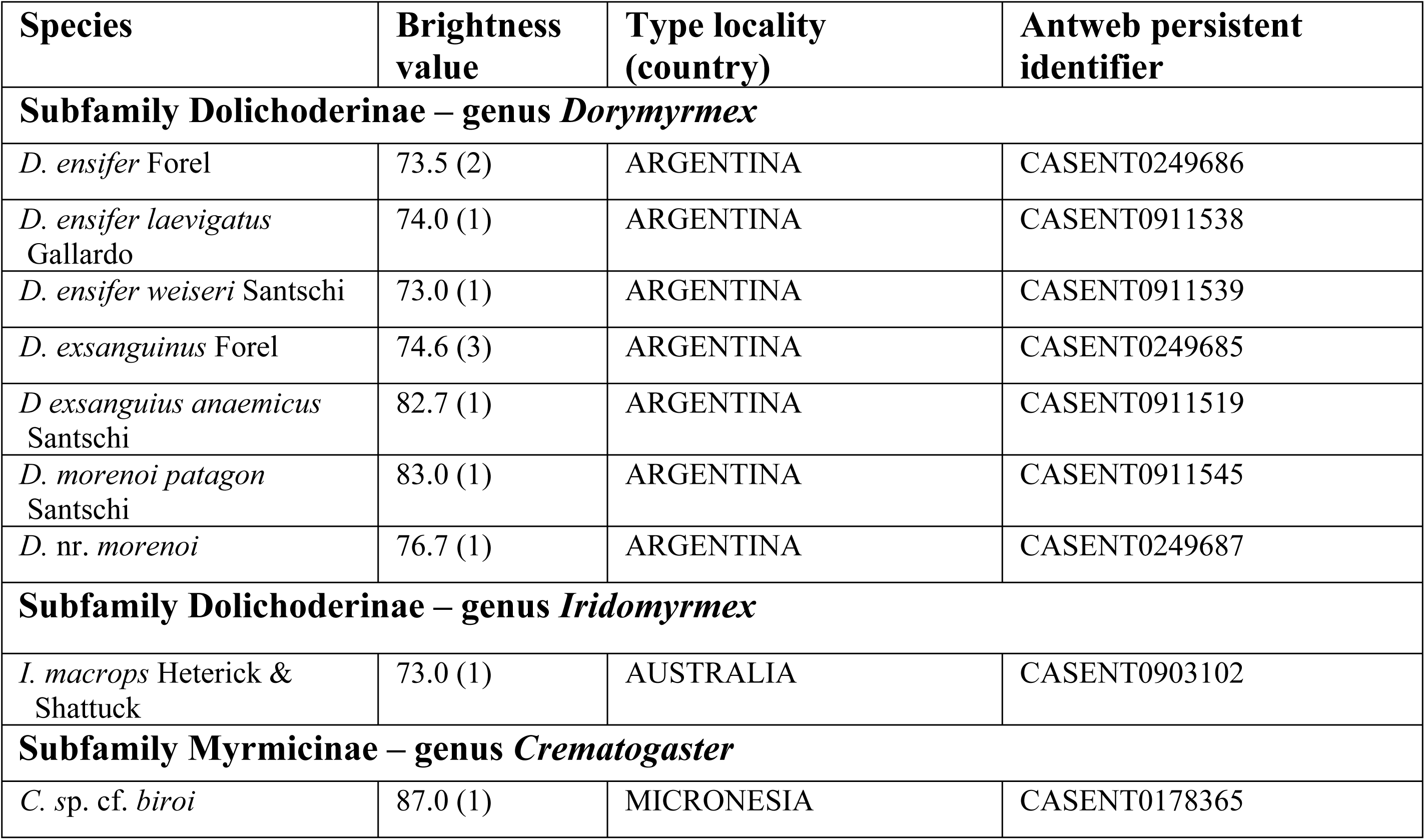
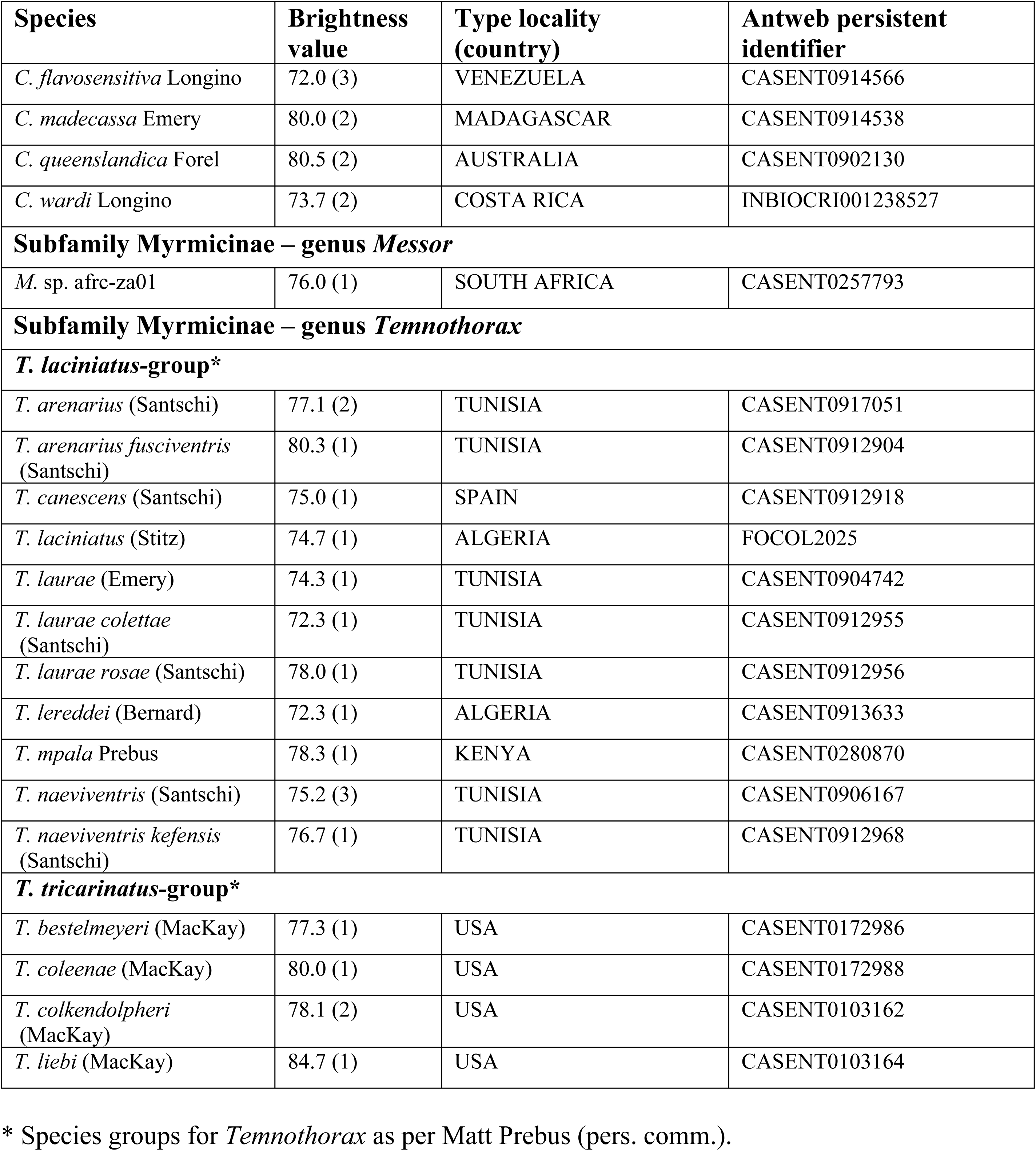
Additional pale ant species (see text) with enlarged eyes in five genera based on photographs examined on Antweb (www.antweb.org). Species are listed alphabetically by subfamily, genus, species group, species, and subspecies. High resolution photographs of each species can be viewed by going to https://www.antweb.org/advSearch.do, then placing the persistent identifier for each taxon in the basic search box. To compare eye size across all species in a genus, place the genus name in the basic search box, click on images, then scroll through the species. Brightness values are given as mean (*n*) (see text).

## Discussion

### Foraging and cuticular color

Worker color was correlated with foraging time across all four genera of ants, suggesting that pale coloration is linked to nocturnal foraging in these and other ants. Alternatively, dark species usually forage diurnally, but some species also forage nocturnally during warm seasons, and several species are largely matinal-crepuscular-nocturnal foragers. Moreover, pale color involves repeated evolution of similar color phenotypes in response to living in dim light to lightless environments both in these ants and in numerous other organisms [13, 14, 24], but it is not a necessary phenotype given the numerous taxa living in similarly dim conditions that have retained their pigmentation.

A species-level phylogeny is available for all four genera such that we can infer the direction of trait evolution. These phylogenies infer that pale color is a derived trait in *Aphaenogaster* [44], *Temnothorax* [47], and *Veromessor* [62], i.e., all most recent common ancestors of pale species were dark, but that it is an ancestral trait in *Myrmecocystus* [43], i.e., pale color was a basal trait in this genus and that these species gave rise to dark congeners. Van Elst [43] also determined that the subgenus *Myrmecocystus* (all pale species) and the genus *Myrmecocytus* as a whole most likely originated from a nocturnal ancestor. Interestingly, the sister genus, *Lasius* [43], contains numerous pale species that are largely subterranean with very small eyes [63, 64].

### Compound eye morphology

Using pale body color as an indicator of nocturnal activity in ants demonstrated consistent correlated adaptations in eye structure across four genera of ants in two subfamilies. When controlled for body size, pale species exhibited convergent morphology for some characters, but not for others: all pale species (except *V. lariversi*) had larger eyes, a larger *D*, and a larger visual span compared to their dark congeners. *Aphaenogaster megommata* and *V.* RAJ-*pseu* also possessed more eye facets than their dark congeners. Alternatively, Δ*ϕ* and *ρ* displayed variable patterns both within and among genera. These general patterns suggest selection on pale species to maximize sensitivity, which is the pattern typical for most nocturnal insects with apposition eyes [33, 65].

Sensitivity (light gathering potential) of an eye is a function of four variables that effect photon capture – *D*, rhabdom diameter, rhabdom length, and focal length [see 27]. Facet area [π/4 × *D*^2^] is one of the more important variables that affects light catching potential [33], and consequently can be used to assess differences between pale versus dark species in each genus. This mean difference was highest for *Myrmecocystus* with facet area for pale species about 2.0– 2.1-fold higher than for their paired dark species (calculated using estimated marginal means as [π/4 × *D*^2^_pale_]/[ π/4 × *D*^2^_dark_]; see Table 4), about 1.3–1.5-fold higher for *A. megommata* compared to its dark congeners, and about 1.5–1.7-fold higher for *T.* sp. BCA-5 compared to its dark congeners. Alternatively, for *Veromessor*, facet area was highest in the dark ***V. smithi***. The mean difference was about 1.05–1.10× higher for ***V. smithi*** than for *V. lariversi* and *V*. RAJ-*pseu*; all other dark congeners had a smaller *D*. This study examined only *D*, suggesting that these sensitivity values are minimum differences between pale and dark species. Pale species of *Myrmecocytus* also differed in that their eyes were more protruding and dome-shaped compared to the more flattened eyes of their dark congeners (Fig 1). These more bulging, dome-shaped eyes result in a greater radius of curvature and possibly a greater visual span field, as well as space for more facets within a given eye area.

Interestingly, the two pale sister species of *Veromessor*, *V.* RAJ-*pseu* and *V. lariversi*, displayed different patterns of eye structure, with *V.* RAJ-*pseu* having larger eyes and more facets than *V. lariversi* (Fig 9). Additionally, eyes of the dark species ***V. smithi*** were smaller with fewer facets than *V.* RAJ-*pseu*, but they were larger with larger *D*’s compared to *V. lariversi*. This may result from the fact that ***V. smithi*** is the most nocturnally-active of all dark species in the genus. One difference between *V. lariversi* and *V.* RAJ-*pseu* and other pale species examined herein is their more yellowish-amber to yellowish-orange color and lower B value, indicating that they are less pale than pale species in the other three genera (see Table 1; Figs 1-4). Differences in eye structure across genera along with variation across species of *Veromessor* are similar to the wide variation in degree of pigment loss and eye degeneration found among cave-dwelling species that is caused by differences in divergence time and intensity of selection [16]. Similar variation occurs for nocturnal foraging bees in which many species have a relatively pale body color, and many but not all species have enlarged compound eyes and ocelli [18].

All of our species had relatively large Δ*ϕ*’s that ranged from 3.5–7°. Pale and dark species varied in their patterns of Δ*ϕ* which were significantly larger in the dark *Temnothorax* and *Aphaenogaster*, which had small eyes with fewer facets, but was larger for the pale *Myrmecocystus* which had numerous eye facets. Moreover, Δ*ϕ* did not decrease for pale species indicating that daily activity patterns have had little effect on the evolution of resolving power.

The eye parameter (*ρ*) measures the tradeoff between sensitivity and resolution, with eyes that require higher sensitivity having larger *ρ* values. Consequently, insects active during high light conditions usually have low *ρ* values that enhance resolution, whereas species active in low light have higher *ρ* values that often exceed 2 um rad [34]. Across our four genera, *ρ* was significantly higher only for the pale *M. navajo* compared to the dark ***M. kennedyi***, with *ρ* for the former species approaching 2 (Fig 11). The higher *ρ* value for *M. navajo* resulted from the combination of significantly larger facets and a significantly larger Δ*ϕ* (Figs 5 & 10). In contrast, ***M. kennedyi*** was the only strictly diurnal forager among all dark species (Table 1), and correspondingly it had the lowest mean *ρ* value (0.91) among all species (Fig 11). The *ρ* value was similar for the other three pairs of congeners, with the dark ***A. occidentalis*** having the highest *ρ* value (2.12) of all species (Fig 10). The lack of significant and consistent patterns across the latter three genera likely reflect the wide range of light conditions under which dark species forage including nocturnal foraging in some seasons (Table 1).

The visual field was larger, usually significantly so, for pale species in all four genera (Fig 13). Moreover, there was no indication that these species had binocular vision in the anterior-posterior direction, that is, they cannot use their eyes for binocular depth perception. This infers that these ants do not use vision to find or capture food items, which aligns with diets that include stationary objects such as seeds, dead insects, and extrafloral nectaries. Instead, it seems likely their eyes are used for detection and orientation relative to land-based and celestial cues used in navigation (see below). In addition, our finding that visual field usually correlated with body size (positively or negatively, depending on the species), contrasted with the pattern for *Cataglyphis bicolor*, in which visual field was independent of body size [66].

Regional variation in *D* is common in insects [67], with these size differences probably related to the different selection pressures on eye structure in each region. Larger facets imply that insects have better vision from regions containing larger facets. In this study, ventral facets were significantly larger in three of the four examined species (along with anterior facets in two species), and they also were largest in the fourth species (***M. kennedyi***) but the difference was not significant (Fig 12). This general pattern suggests that ventral facets are important for vision in both diurnal and nocturnal activity, perhaps as a mechanism for optic flow to measure distance [see 68, 69].

### Other pale ants with enlarged eyes

Our survey of images on Antweb (https://www.antweb.org/) revealed numerous additional species of pale ants with enlarged eyes in the clades studied here and in other genera. Moreover, these coupled traits appear to have evolved independently multiple times in multiple genera across at least three subfamilies. Numerous additional pale species undoubtedly occur given the limited scope of museum specimens available, and it is likely that many pale species remain to be discovered. As these are located, our technique for measuring brightness provides a tool for mapping patterns of pigment loss within and across ant genera.

One commonality among pale species examined herein and in Table 6 is that many of these species largely are restricted to desert and semi-arid habitats. As such, these species possess visual adaptations to be nocturnal specialists in extreme environments in a manner similar to heat tolerance adaptations possessed by their thermophilic diurnal counterparts such as *Myrmecocytus kennedyi* and *Forelius* spp. [28; R.A. Johnson, pers. obs.] in the New World, and *Cataglyphis* spp. and *Melaphorus bagoti* in the Old World [70, 71]. The open, exposed nature of their foraging environment lacks overstory which suggests that these species can obtain navigation cues from local landmarks via their enlarged eye facets, but probably only horizon and lunar night sky cues. However, at this point, nothing is known about navigation in any pale species, and among dark species, orientation and navigation have been examined only in the column-foraging, mostly diurnal ***V. pergandei*** [72–74]. There is much to learn about how ants use their eyes both at night and during the day.

### Ocelli

Size of the anterior ocellus varied among pale and dark species of *Mymecocystus*. In larger species, the anterior ocellus was smaller in pale compared to dark species, but this difference largely disappeared for smaller species (Fig 6). The two largest pale species (*M. mexicanus*-01, *M. mexicanus*-02) also displayed size-dependent presence of the anterior ocellus as it was present only in larger workers. The anterior ocellus also was absent in nearly all workers of the intermediate sized *M. navajo*. The pattern was mixed for smaller species because the anterior ocellus was largest for the dark ***M. yuma***, intermediate for the dark ***M. kennedyi*** and pale *M. ewarti*, and smallest for the pale *M. testaceus* (Fig 6). In contrast, the anterior ocellus is typically larger in nocturnal compared to crepuscular and diurnal flying bees and ants [31, 75–78], as well as in pedestrian workers in the ant genus *Myrmecia* [79].

Absence of the anterior ocellus in some to most workers of some pale species displays a phylogenetic component. Pale species in which some to most workers lacked the anterior ocellus fell into one clade, while all other species that always have an anterior ocellus were in two other clades [see 43]. We were unable to examine the two other pale species (*M. pyramicus*, *M. melanoticus*) because specimens were unavailable, but this phylogenetic association predicts that the anterior ocellus always is present in *M. pyramicus* and that it is only present in larger workers of *M. melanoticus* (see Fig 6). To our knowledge, these are the only known ant species in which workers display intraspecific variation in presence-absence of the anterior ocellus, making them excellent candidates to examine evolution, development, and function of the anterior ocellus, as well as how such variation affects forager orientation and navigation (see below). Foraging behavior is poorly documented in pale species, but it appears that both small and large workers of *M. mexicanus*-02, i.e., those with and without an anterior ocellus, leave the nest to forage (J. Conway, pers. comm.).

The function of ocelli in ant workers is poorly understood because most species lack ocelli (notable exceptions include the genera *Cataglyphis*, *Formica*, *Myrmecocystus*, *Polyergus* in the subfamily Formicinae; *Myrmecia* in the subfamily Myrmeciinae). However, pedestrian workers that have ocelli provide a functional contrast to that of conspecific flying queens and males, where ocelli are almost always present. Flying insects have three ocelli that serve the general purpose of sensing polarized light for navigation and maintaining flight stability, whereas workers use their ocelli to detect polarized light for navigation in *Cataglyphis bicolor* [80] or to gather light in *Myrmecia* [79].

Lastly, compound eyes and ocelli provide separate and functionally different visual pathways, so it is instructive to examine for convergence in the two pathways. Two studies compare compound eyes and ocelli between nocturnal and diurnal ant species. In leafcutter ants (genus *Atta*), both the ocelli and eye facets were larger in nocturnal compared to diurnal species of both flying queens and males, while eye area was similar for species in both activity groups [76]. The other study examined workers of four species of *Myrmecia* also finding that both the ocelli and eye facets were larger in nocturnal compared to diurnal congeners, while number of facets was similar for species in both activity groups [27, 79]. Alternatively, this study found that pale species had an anterior ocellus that was similar in size to smaller than comparable dark congeners, but that eye facet diameter and eye size were larger for pale compared to dark species of *Myrmecocystus*; facet number was similar for species in both activity groups. Moreover, facet size and eye size/number of facets display similar patterns across these studies, whereas relative size of the ocelli varied across genera.

## Conclusions

This study provides a first overview of variation in external eye structure across several ant genera that compares closely related pale and dark congeners. Our observations on body coloration and eye structure allow several statements about their visual ecology. First, the correlation between ant body color and activity period parallels that found in other animals. The specific selective factors shaping this correlation await more detailed work on the costs and benefits of cuticular pigmentation. Second, pale, above ground foraging ants have enlarged rather than reduced or no eyes relative to their dark congeners, suggesting that vision is important for both nocturnal and diurnal species across several lineages. That pale species possess optical adaptations to maximize sensitivity over resolution, which is the pattern typical for most nocturnal insects with apposition eyes [33, 65], also suggests that vision plays a role in navigation for these nocturnal ants. Third, the visual field span and mild regional variation in *D* suggest that their eyes are not adapted to gather detailed visual information from any specific region in the space around the ant, but rather they are gathering relatively low quality information from a large part of the space around them. Fourth, the mild differences in eye structure between pale and dark species suggest both groups use their eyes in similar ways, and they are consistent with observations that these ants use their vision in navigation guided by celestial and large landmark cues. Field studies that detail foraging behavior and navigational skills would complement these data. Additional research should be done to more thoroughly determine optical sensitivity. This study only examined facet diameter, but data are needed on rhabdom diameter, rhabdom length, focal length, and neural adaptations to more completely determine and compare optical sensitivity [see 27]. We also note that activity period is the primary difference between our pale and dark species given that life history and behavior are similar for species within each genus, i.e., most species are solitary foragers that harvest seeds, or scavenge for debris, dead insects, and plant exudates [28, 48: R.A. Johnson, pers. obs.], probably using olfactory and/or tactile cues.

Of the genera examined herein, we believe *Myrmecocystus* has the most potential for further study given the consistently large variation in eye structure between pale and dark species (eye area, *D*, *ρ*, visual span), combined with the fact that most species are strongly polymorphic such that traits can be compared allometrically [see 66]. Additionally, this is the only known genus with pale species that possess ocelli, such that it provides an excellent group to examine internal eye structure and to compare evolution of both eyes and ocelli. The flying queens and males might also be examined for comparative study of the sexual castes, especially given that the queen of *M. navajo* has extremely large ocelli.

## Acknowledgments

We thank Brian Fisher, Michele Esposito, and Antweb for the high resolution photographs, Christian Rabeling for use of his microscope and photographic programs, and Matt Prebus for the loan of specimens.

## Supporting information

All data will be deposited in the Dryad Digital Repository.

## Author contributions

Conceptualization: Robert A. Johnson.

Data curation: Robert A. Johnson, Ronald L. Rutowski.

Formal analysis: Robert A. Johnson, Ronald L. Rutowski.

Methodology: Robert A. Johnson, Ronald L. Rutowski.

Writing - original draft: Robert A. Johnson.

